# Hybridogenesis as an intermediate step between sexual reproduction and parthenogenesis

**DOI:** 10.1101/2025.09.19.677091

**Authors:** Alexander Brandt, Guillaume Lavanchy, Vincent Mérel, Luca Soldini, Morgane Massy, Zoé Dumas, Emelyne Gaudichau, Marjorie Labédan, Falon Pasquier Genoud, Marc Bastardot, Tanja Schwander

## Abstract

Many organisms reproduce through non-canonical modes such as parthenogenesis or hybridogenesis (clonal transmission of one parent’s chromosomes), but whether these arise abruptly or stepwise from each other remains unclear. We address this in the stick-insect genus *Bacillus*, which harbors several hybrid lineages with diverse reproductive modes. From haplotype-resolved phylogenies of >500 wild-caught individuals, we infer a single, recent (∼8,000 years) origin of all hybrids. The ancestral hybrid reproduced via hybridogenesis, which subsequently diversified into parthenogenesis and, twice independently, into triploid lineages. Laboratory crosses recapitulate this trajectory, where each step facilitated the next. These findings reveal how a single genomic perturbation can act as a catalyst for evolutionary innovation, turning the loss of sex into a driver of diversification rather than a dead end.

## Introduction

The ability to reproduce is a fundamental property of life. Most animals reproduce through sex, which involves producing gametes via meiosis and fusing them with those from another individual (*1*). Modifications to these key processes, however, have resulted in the repeated evolution of alternative reproductive modes across many taxa (*2*). This includes parthenogenesis (reproduction without fertilization, which is often clonal) and hybridogenesis, whereby only one set of chromosomes (haploset), usually the maternal one, is transmitted clonally to the offspring, and the other discarded (a.k.a. hemiclonal reproduction; (*3*)). While considerable research has focussed on the costs and benefits of these alternative reproductive modes (*4*, *5*), how they originated in the first place remains largely enigmatic (*6*, *7*).

Alternative reproductive modes require several modifications to canonical meiosis, imposing strong constraints on their evolution (*8*). Some taxa are much more prone than others to evolve alternative reproductive modes (e.g. (*9–11*)), suggesting that they might be characterized by fewer constraints, thus increasing the frequency of transitions. This could explain the repeated evolution of these alternative reproductive modes in some taxa (*9*, *12–14*) as well as their frequent co-occurrence in the same clades (reviewed in (*15*)). An alternative explanation is that initial transitions from sexual reproduction to alternative reproductive modes can themselves promote further diversification. Such diversification could notably be triggered by exacerbated genetic conflict, as these modes often involve non-Mendelian transmission creating new opportunities for parental chromosomes to increase their own transmission.

Because reproductive mode transitions can be rapid and intermediate states transient, only the end products are usually visible in nature. Studying transitions between reproductive modes thus relies either on the ability to induce new reproductive modes via laboratory crosses (*16–18*), or on serendipitous findings of intermediate states (*19*). The stick insect genus *Bacillus*, which features a remarkable diversity of reproductive modes (summarized in Figure 1), provides unique opportunities to study transitions between alternative reproductive modes.

**Figure 1:**
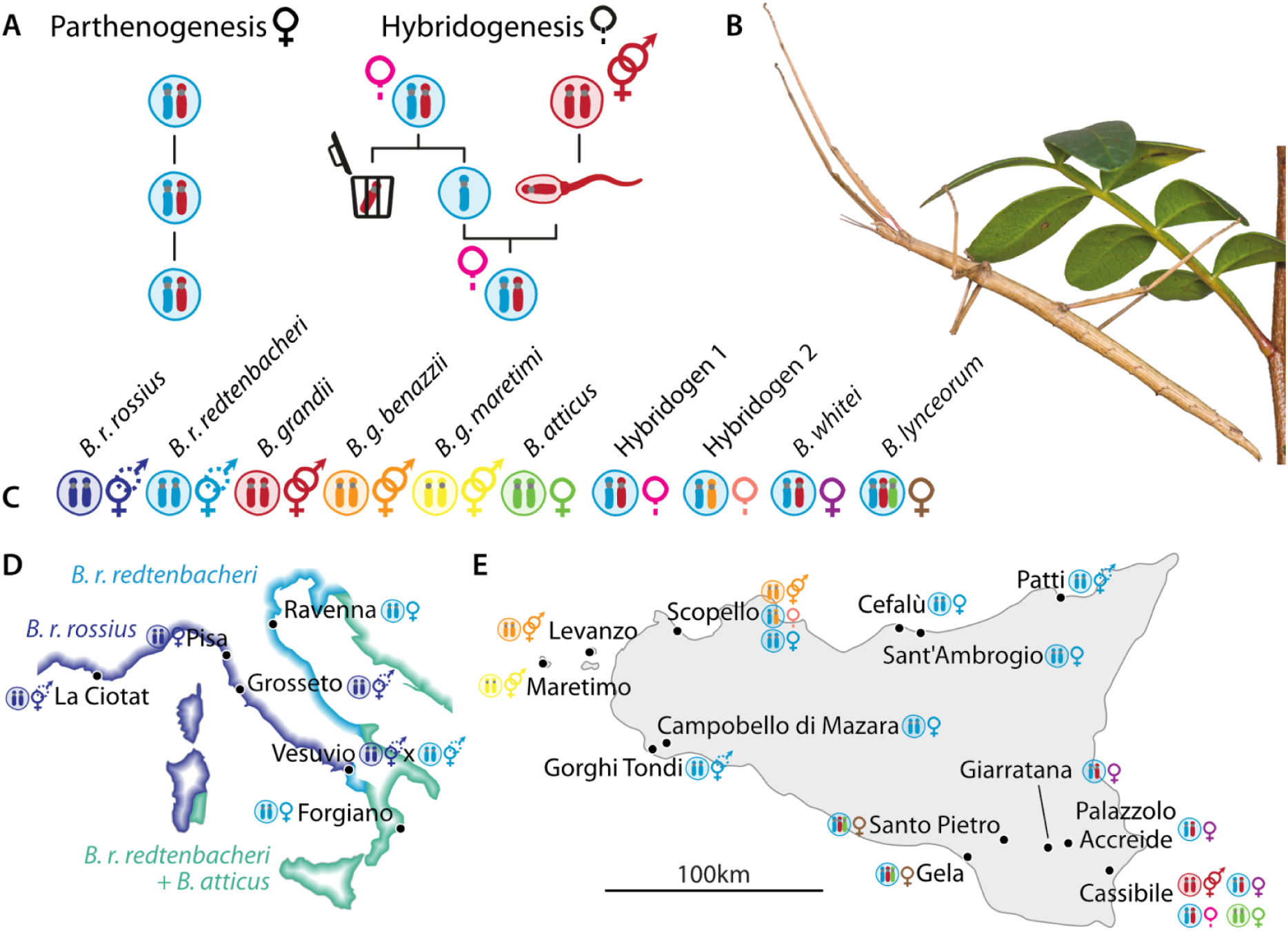
Overview of the *Bacillus* system. **A:** Schematic overview of parthenogenesis and hybridogenesis. **B**: An adult female *Bacillus* on one of her host plants, the lentisk *Pistacia lentiscus*. **C:** Genomic composition and reproductive modes of the different lineages studied. One species, *B. grandii*, is obligately sexual. There are three known subspecies (*B. g. grandii*, *B. g. benazzii* and *B. g. maretimi*). A second one, *B. rossius*, is facultatively parthenogenetic, with all-female and bisexual populations. A third species, *B. atticus*, reproduces via obligate female-producing parthenogenesis (although rare male production has been documented (*20*)). Two lineages are F_1_ hybrids between the subspecies *B. r. redtenbacheri* of *B. rossius* and *B. g. grandii*. The first one (*B. whitei*) reproduces by obligate parthenogenesis. The second one (hybridogen 1) reproduces by hybridogenesis, whereby the *B. rossius* haploset (haploid set of chromosomes) is transmitted clonally, while the *B. g. grandii* haploset is eliminated. The F_1_ constitution is then restored by mating with *B. g. grandii*. A second hybridogenetic lineage exists, which differs from the first one in having a haploset from *B. g. benazzii* instead of *B. g. grandii* (hybridogen 2). The two hybridogenetic lineages occasionally reproduce through androgenesis, whereby both haplosets are eliminated from the egg and the progeny ends up with two copies of the paternal *B. grandii* genome. Finally, an allotriploid asexual, *B. lynceorum*, has one chromosome set from each of *B. r. redtenbacheri*, *B. g. grandii* and *B. atticus*. All hybrid lineages have *B. r. redtenbacheri* as maternal ancestor (*21*). **D:** Geographic distribution of *Bacillus* lineages and location of sampling sites in Italy and the surrounding Mediterranean basin. Estimated coastal distribution ranges are colored according to species following (*21*, *22*). **E**. Map of Sicily, with species found in each sampling location.

Here, we investigate whether *Bacillus* hybrids with different reproductive modes evolved independently or share a single hybrid origin, followed by reproductive mode diversification. To this end, we generate reference genome assemblies or pseudo-references for all parental lineages and collect genomic data for over 500 wild-caught stick insects, enabling us to separately reconstruct the evolutionary histories of the maternal and paternal genomes of the parthenogenetic and hybridogenetic hybrids. Phylogenetic topology tests combined with estimates of divergence allow us to resolve the origins of these hybrids and infer transitions between hybridogenesis and parthenogenesis and between diploid and triploid lineages. Finally, we test the plausibility of the inferred transitions by recapitulating them through experimental crosses of extant lineages.

## Results and Discussion

We sampled natural stick insect populations from Sicily (13 populations) and mainland Italy (6 populations). We compiled RAD sequencing data for 504 individuals (108 new and 396 previously published (*23*)) representative of all lineages. Because morphology-based identification of *Bacillus* stick insects is error-prone, we genetically identified all individuals based on genetic similarity to previously characterized reference individuals (PCA; see Suppl. Results 1; Figure S1; Methods; (*23*)). We phased the reads of 172 hybrid individuals by competitive mapping, i.e. simultaneously mapping the reads of each individual to the assemblies/pseudoreferences of all of its parental species (see Suppl. Results 2 & 3; Figures S2 & S3). Calling SNPs separately with the reads that mapped to each assembly then enabled us to reconstruct haploset-specific genotypes and phylogenies of *B. rossius* (maternal ancestor; Figure 2A; Suppl. Results 4; Figure S4) and *B. grandii* (paternal ancestor; Figure 2B; Suppl. Results 5; Figure S5).

**Figure 2.**
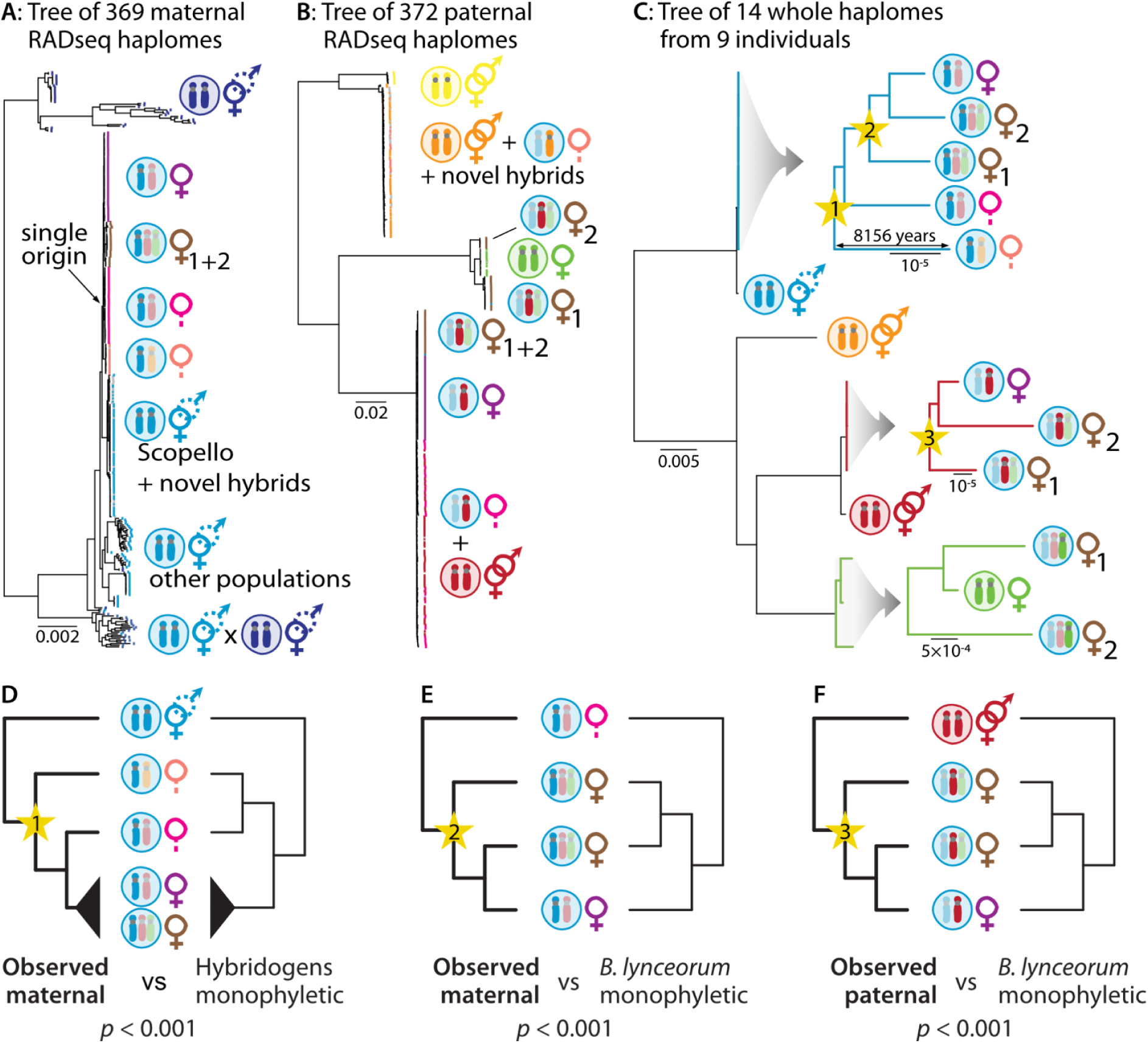
Haplome-specific phylogenies for **A:** haplomes of *B. rossius* origin (maternal ancestor) and **B:** haplomes of *B. grandii* origin (paternal ancestor) from the RADseq data. Colors at the tips = lineage, colors as in Figure 1. Novel hybrids refer to *B. rossius* - *B. g. benazzii* F_1_ hybrids stemming from ongoing hybridization between sexual lineages and whose reproductive mode is unknown. **C:** Whole-haplome phylogenies. The three smaller trees to the right correspond to zoomed subsets of the main tree. Yellow stars designate nodes at which topology tests were performed. **D - F**: Topology tests. The topology in bold is significantly better supported.

All hybrid lineages formed a monophyletic lineage in the maternal *B. rossius* phylogeny, indicating a single origin followed by diversification via transitions between alternative reproductive modes (Figure 2A). Independent origins via several hybridization events would result in some lineages of *B. rossius* clustering with a subset of the hybrids. While the existence of such lineages (which may now be extinct) can never be formally excluded, our dense sampling within and beyond Sicily (Figure 1D, E) ensures broad coverage of the extant diversity. All hybrids were most closely related to *B. r. redtenbacheri*, the subspecies that is currently present in Sicily, confirming their Sicilian origin.

All hybrid lineages seemed to have originated over a short period, as indicated by very short branch lengths. In order to improve resolution of the evolutionary relationships among hybrid lineages, we built an additional phased phylogeny of 14 haplomes from whole-genome data of nine individuals (8 new + 1 published (*23*)) representing all four parental species plus one representative of each the four expected hybrid lineages (hybridogen 1, hybridogen 2, *B. whitei* and *B. lynceorum*; Suppl. Results 6). We included a fifth hybrid lineage as we discovered two distinct triploid *B. lynceorum* lineages in the paternal phylogeny (Figure 2B). Note that we also discovered several F_1_ hybrid individuals that did not cluster into lineages and which likely represented novel hybrids stemming from contemporary interspecific matings between parental species (see below). These were not included in the whole genome analyses as their reproductive mode is unknown.

The whole-genome phylogeny confirmed the single origin of all hybrids (Figure 2C) and indicated that the original hybridization event between *B. rossius* and *B. grandii* occurred over ∼8000 generations (∼8000 years) ago and resulted in a hybridogenetic hybrid (Fig 3A). This was indicated by the independent (paraphyletic) and basal position of the two hybridogenetic lineages, and longer branch lengths of hybridogens. Whether hybridization triggered hybridogenesis is not certain, as we cannot exclude that hybriodogenesis-like reproduction already existed in specific *B. rossius* lineages.

**Figure 3.**
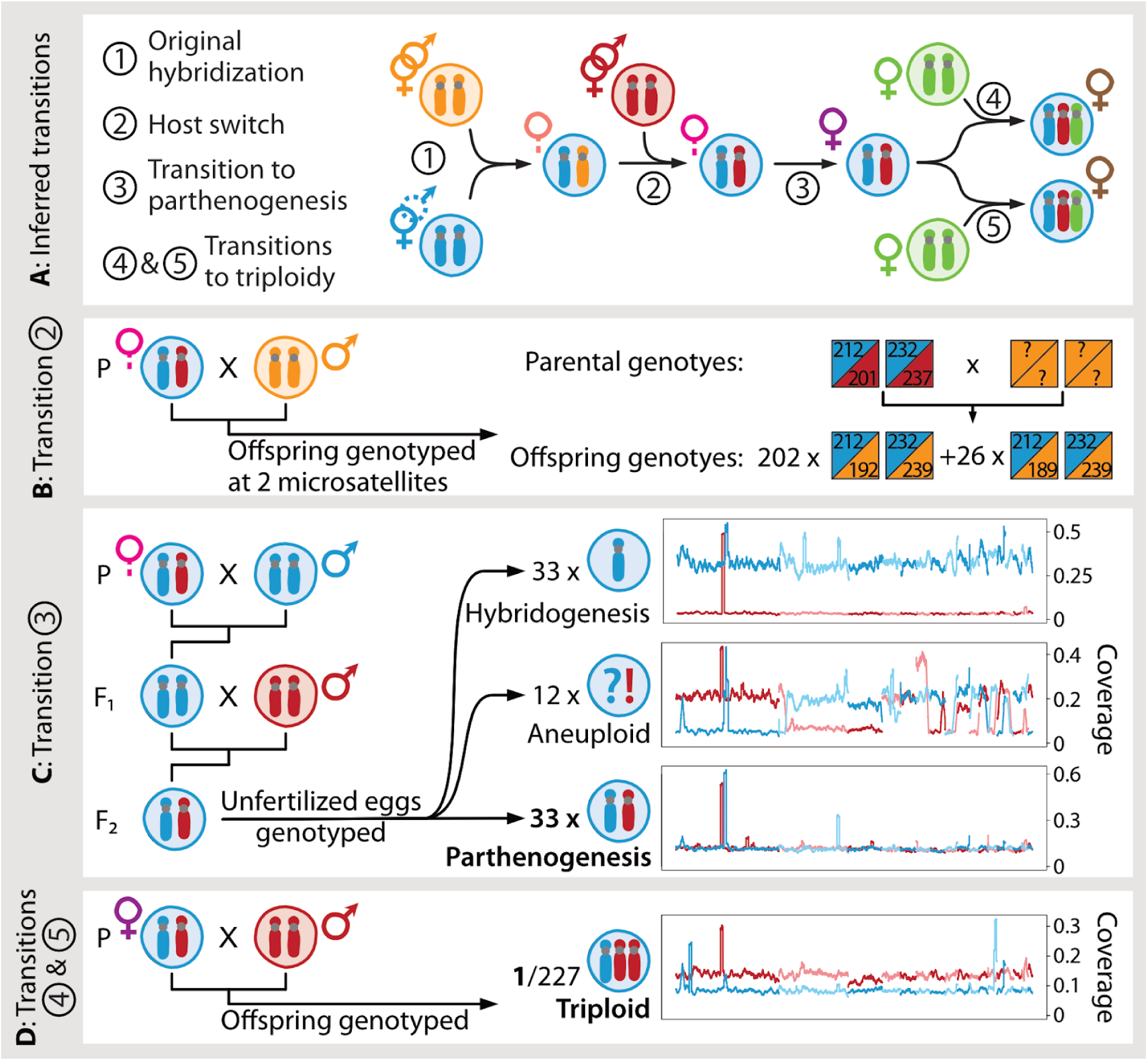
**A**: summary of the most likely transitions between reproductive modes in the different lineages of *Bacillus* inferred from our phylogenetic analyses. Colors as in Figure 1. **B-D**: experimental evidence for the main transitions. **B**: genotypes of the mother and 228 female hatchlings at two microsatellite loci (two boxes) showing the replacement of the *B. g. grandii* allele (red) with the paternal *B. g. benazzii* allele (orange). **C**: Spontaneous transition to parthenogenesis in some individuals with admixed *rossius* haplome. Right: Average coverage of the *B. grandii* (red) and *B. rossius* (blue) haplomes in 100kb windows along the genome obtained by competitive mapping of RADseq reads, for three unfertilized eggs representative of the different observed patterns. Alternating shades represent different chromosomes. **D**: Transition to triploidy by incorporation of an additional *B. g. grandii* genome in an egg of *B. whitei* fertilized by *B. g. grandii*. Right: Average coverage of each genome in 100kb windows along the genome obtained by competitive mapping of RADseq reads. Coverage of the *B. grandii* genome is ∼2x that of *B. rossius*.

Hybridogen 2 (*B. rossius* x *B. g. benazzii*) appears to be the oldest hybrid lineage, as it formed the outgroup to all other hybrids (Figure 2C) and clustered with *B. r. redtenbacheri* on the mitochondrial tree, both lacking a substitution that was common to all other hybrids (Suppl. Results 6; Figure S6). Paraphyly of hybridogens (with hybridogen 1 more closely related to parthenogenetic hybrids) was better supported than an alternative topology where hybridogens were constrained to be monophyletic (Shimodaira-Hasegawa test, *p* < 0.001; Figure 2D). The second hybridogenetic lineage (hybridogen 1) likely emerged following a host switch, i.e. by replacing the *B. g. benazzii* haplome with one of *B. g. grandii* upon mating with that species. Note that a scenario where the original hybridization happened between *B. g. grandii* and *B. rossius*, followed by a host switch to *B. g. benazzii* after the split with hybridogen 1, is equally parsimonious. Our experimental crosses provided experimental evidence that host switches can happen easily (host switch occurred in all 228 offspring from 12 crosses genotyped; Suppl. Results 7; Figure 3B), mirroring previous findings (e.g. (*24*)). Host switches in hybridogens were also described in wild *Hexagrammos* fish (*25*).

All parthenogens then derived from the ancestor of hybridogen 1, with which they share haplosets from both *B. r. redtenbacheri* and *B. g. grandii*. This means that the transition from hybridogenesis to parthenogenesis did not require entirely new haplosets to enter into contact. Rather, the transition to parthenogenesis after generations of hybridogenesis suggests a shared proximate basis of these two reproductive modes. Notably, both reproductive modes probably rely on premeiotic duplication of the genome (the maternal haplome in hybridogens and the whole genome in parthenogens (*26*)). Pre-existing duplication in hybridogens may thus have facilitated the transition to parthenogenesis. The evolution of parthenogenesis from hybridogenesis further requires the transmission (instead of elimination) of the paternal genome, resulting in the production of unreduced F_1_ hybrid eggs. We were able to experimentally induce such production of unreduced F_1_ hybrid eggs by introgression of sexual *B. rossius* alleles into the hybridogenetic haplome via experimental crosses (Suppl. Results 7; Figure 3C).

Our whole-genome tree also supported the paraphyly of the triploid *B. lynceorum*, with one lineage (hereafter *B. lynceorum* 2) forming a monophyletic group with *B. whitei* and the other (*B. lynceorum* 1) being their outgroup, both on the maternal *B. rossius* and paternal *B. grandii* phylogenies (Figure 2C, E, F). The *B. atticus* haplomes of the two *B. lynceorum* lineages were also diverged, further supporting separate origins. Their phylogenetic position indicates that *B. lynceorum* 2 likely originated from mating between a female *B. whitei* and a rarely-produced male of *B. atticus* (*20*). *B. lynceorum* 1 could have originated from a cross between either *B. whitei* or a female hybridogen 1 and a male *B. atticus*, in which case it would represent a second transition from hybridogenesis to parthenogenesis, this time associated with hybridization. Although alternative scenarios (e.g. *B. whitei* originating from the loss of the *atticus* haplome from *B. lynceorum*) cannot be formally excluded, a transition from *B. whitei* appears more likely. Secondary ploidy increases in parthenogenetic species are reported in other taxa (e.g. (*27–29*)), and specifically, the topologies of the *rossius* and *grandii* haplomes in *B. whitei* and both *B. lynceorum* lineages perfectly mirror each other, which is expected under clonality but not under hybridogenesis, where a different *grandii* genotype could have been incorporated. Additional support for our favored scenario comes from our ability to replicate the diploidy to triploidy transition experimentally. We crossed female *B. whitei* with *B. grandii* males (since accidental *B. atticus* males are extremely rare) and found that one out of 227 offspring was triploid with two *B. grandii* haplosets (Figure 3D), corroborating findings by (*30*).

The relationship between asexuality and polyploidy has long been noted (*31*) and triploidy is often assumed to cause transitions to asexuality, but a causal relationship is rarely shown (*6*). Here, the transition to triploidy was facilitated by parthenogenesis via endoduplication (which is the proximate mechanism of parthenogenesis in both *B. whitei* and *B. lynceorum* (*26*, *32*), since segregation between copies of the same chromosome releases the constraint that chromosomes should be present in pairs for segregation to occur properly during regular meiosis. Triploidy was thus a consequence, and not a cause, of parthenogenesis. Interestingly, the third ancestor of *B. lynceorum*, *B. atticus*, is also parthenogenetic. However, the presence of *B. rossius* mitochondria in *B. lynceorum* (*21*, *23*) indicates that the *B. atticus* haploset was transmitted by a male. Such rare, “accidental” males are sometimes found in asexual species, especially with XX/X0 sex determination where they can easily arise by X aneuploidy, and with a documented example in *B. atticus* (*20*).

Finally, two elements in our results support the idea that only a subset of interspecific genome combinations can lead to asexuality: the fact that parthenogenesis arose only once in ∼ 8000 years in two different hybridogenetic lineages, as well as the serendipitous finding of previously undescribed hybrid individuals in our RADseq screen of wild-caught stick insects. 20 F_1_ hybrids (“novel hybrids” in Figure 2) clustered with pure *B. r. redtenbacheri* from the same area on the maternal tree (Figure 2A; Figure S4). This suggests that they are newly-formed F_1_ hybrids between *B. r. redtenbacheri* and *B. g. benazzii*. Although their fertility and reproductive mode was not assessed, available evidence suggests that they are most likely sterile. The presence of a large proportion of males (12 out of 20) and the fact that they were not monophyletic excludes the possibility that they are a separate hybridogenetic or parthenogenetic lineage. In addition, the absence of backcrosses or F_2_ genotypes points to hybrid sterility. These findings are in line with the Balance hypothesis (*33*), according to which asexual taxa emerge from crosses between lineages that have diverged enough to disrupt normal meiosis, but not enough to result in too many incompatibilities, as was observed in other taxa (*16*, *18*, *34–37*).

In summary, reconstructing haplotype-specific phylogenies for *Bacillus* hybrids with alternative reproductive modes and their parental species enabled us to infer the number and direction of transitions between reproductive modes (summarized in Figure 3). A single original hybridization event between the maternal (*B. rossius*) and paternal (*B. grandii*) sexual ancestors gave rise to a hybridogenetic hybrid lineage. A second hybridogenetic lineage arose by host switch, i.e. by substitution of the paternal genome with that of another subspecies of *B. grandii*. Obligate parthenogenesis then evolved from hybridogenesis, in what is the first documented case of such a transition that we are aware of. Hybridogenesis probably facilitated this transition by combining a new set of paternal chromosomes with the maternal chromosomes every generation, thus creating repeated opportunities for compatible genome combinations to emerge. Finally, two independent transitions to triploidy happened from the diploid parthenogenetic lineage. This was facilitated by premeiotic endoduplication, which releases the constraint on homologous chromosome pairing during meiosis. Overall, our results indicate that the diversity of reproductive modes in *Bacillus* stick insects was achieved via stepwise asexual diversification rather than repeated transitions from sexual ancestors. Each transition released some of the constraints hampering the evolution of unorthodox reproductive modes, thus facilitating the next one. The diversification of reproductive modes in *Bacillus* reveals how a single genomic perturbation can act as a catalyst for evolutionary innovation, turning the loss of canonical sex into a driver of diversification rather than a dead end.

### Data Availability

The reference genome is available at NCBI under BioProject PRJNA1251886, the whole genome data of eight additional lineages (including males used for the introgression experiment) is available under BioProject PRJNA1255460, the demultiplexed RADseq reads (field collected individuals and crosses) are available under BioProject PRJNA1031135, and all scripts are available on Github (https://github.com/AlexPopalex/tree).

## Acknowledgements

We would like to thank our many helpers taking care of the stick insects and supporting field and lab work over the last five years: Astrid Anaí Oliva De León, Coline Boulard, Alexandre Roland, Simon Vogel, Sophie Dupertuis, Arno Gattoliat, Jérôme Smyrliadis, Noah Goël, Emmanuelle Tran, Asia Lahici, Neige Chapuis. We thank Barbara Mantovani, Valerio Scali and colleagues for sharing their knowledge of *Bacillus*. This work was supported by Swiss FNS grant 310030_212232, funding from the University of Lausanne, and funding from the European Union’s Horizon 2020 research and innovation program under the Marie Skłodowska-Curie grant agreement no. 101024860.

## Methods

### Sample collection

We searched for *Bacillus* stick insects in September and October of 2017, 2018, 2020 and 2021 on the island of Sicily and along the coastal ranges of Italy and France by surveying host plants (*Rubus sp.* & *Pistacia lentiscus*) at night. For an overview of sex, sampling year and GPS location per sample, see Table S1 (https://github.com/AlexPopalex/tree/) and Figure 1D,E.

### Wet lab and sequencing

#### Whole-genome data for all lineages

We used PacBio (Pacific Biosciences) Hifi long reads for one individual of each of *B. g. grandii*, *B. whitei* and both *B. lynceorum* lineages, and Illumina short reads for one individual of each of *B. atticus*, *B. g. benazzii*, and for the *rossius* haplotype of both hybridogen 1 and 2. In addition, we also used Illumina short reads for a pool of the four *B. rossius* males that were used in the crosses (see Experimental validation of inferred transitions below).

To generate PacBio HiFi long reads, we ground animals (½ of the body without gut) in liquid nitrogen using a Cryomill (Retsch). We then extracted High Molecular Weight (HMW) DNA with G/20 Genomics Tips by Qiagen according to manufacturer’s instructions. Library preparation (SMRTbell Express Template Prep Kit 2.0) and sequencing (2 SMRT cells on PacBio Sequel II, and 1 SMRT cell on PacBio Revio for *B. lynceorum* Gela4) was carried out by the Lausanne Genomic Technologies Facility, Lausanne University, Switzerland (LGTF).

To generate Illumina short reads, we extracted gDNA with the Qiagen Biosprint 96 workstation following manufacturer protocol from one leg from a single adult (*B. g. benazzii* and *B. atticus*). In order to only sequence the *rossius* haplotype of hybridogen 1 and 2, we extracted DNA from a pool of eggs (see Table S1). For the four *B. rossius* males, DNA was extracted from one leg of each male and then pooled equimolarly and sent to the LGTF for library preparation (Nextera DNA flex) and Illumina sequencing on one lane of an SP flowcell of the NovaSeq (paired-end, 150bp).

#### Scaffolding *B. grandii grandii* assembly

The same individual used for PacBio HiFi long read sequencing was used to generate Hi-C conformation capture reads for scaffolding. After grinding ½ of the body without the gut in liquid nitrogen with a Cryomill, we cross-linked tissues with the Proximo Hi-C kit and sent it to Phase Genomics (Seattle, USA) for sequencing.

#### RADseq of field-caught adults and eggs

We generated ddRAD sequencing reads for 158 individuals and combined them with 396 individuals from (*23*). In addition, we also generated ddRAD sequencing reads for 93 unfertilized eggs used in the crosses for experimental validation of the inferred transitions (see below). We extracted DNA with the Qiagen Biosprint 96 workstation following manufacturer protocol and built double-digest RAD sequencing libraries following the protocol of (*23*). Libraries were sequenced single-end on a MiSeq with 150bp length reads.

### Species determination

For (sub)species determination/corroboration, we compared the genotypes of all new samples (obtained from RADseq or whole genome sequencing [WGS]) to the genotypes of the 397 individuals with known species identity (*23*). We removed adapter sequences and quality-trimmed 3’ ends of RAD and WGS short reads using cutadapt v2.10 and trimgalore v0.6.6 with default parameters (--paired option for WGS short reads; https://github.com/FelixKrueger/TrimGalore (*38*)). We mapped single-end RAD and paired-end WGS short reads to the reference genome of *B. r. redtenbacheri* using bwa mem v0.7.17 with default options (*39*). We mapped PacBio long reads to the reference genome of *B. r. redtenbacheri* using minimap2 v2.14 with the option -x map-hifi and allowing for up to 50 secondary alignments (-N 50 (*40*)). We sorted alignments using samtools and removed unmapped reads and secondary and supplementary alignments as well as alignments with mapping quality below 20 (*41*). For paired-end WGS short reads we only retained properly paired reads. Next, we called variants using bcftools mpileup and bcftools call (*42*) with the option --multiallelic-caller and retaining only variant sites (--variants-only). We filtered variants using bcftools (*42*). We removed duplicate records, indels, mono- and polyallelic sites (keeping only biallelic sites), sites with variant quality < 20 and genotypes with depth < 4. We removed individuals with > 75% missing genotypes (leaving 527 samples). We did pca calculation using plink v1.9 (*43*).

All commands, see “tree_species.sh” (https://github.com/AlexPopalex/tree/).

### Genome assembly

We constructed a *Bacillus grandii grandii* reference genome using a combination of HiFi (∼ 28X) and Hi-C sequencing (∼ 64X). We assembled HiFi reads using hifiasm v0.18.5 (*44*). We then purged putative haplotigs from the assembly as follows: We estimated local coverage by mapping HiFi reads to the assembly using Minimap2 (v2.24, -x map-hifi --secondary=no (*40*)). We investigated homology between contigs of low coverage, i.e. < 18X, using Purge Haplotigs v 1.1.2 (*45*). We filtered out duplicated contigs, as well as contigs of extreme coverage (equal to 0 or superior to 90X). After purging, we performed a decontamination step using blobtools v1.1.1 (*46*) with contigs blasted to the ncbi nt database (v2.12.0, -max_target_seqs 10 -max_hsps 1 -evalue 1e-25 (*47*), (*48*). We filtered out non-metazoan contigs using the script contamination_filtration.py (*49*).

We scaffolded contigs using Hi-C reads. We cleaned the reads with TrimGalore (v0.6.7, -q 30 --nextera --length 100; https://github.com/FelixKrueger/TrimGalore (*38*)), and aligned them to the decontaminated assembly following the Arima-HiC pipeline (https://github.com/ArimaGenomics/mapping_pipeline). We mapped forward and reverse reads separately using bwa v2.12.0 (*50*). We filtered out chimeric reads and combined alignments using Arima scripts and specifying a minimum mapping quality of 57. We removed PCR duplicates using picard tools MarkDuplicates (http://broadinstitute.github.io/picard/). We performed scaffolding using yahs v1.2a.2, -q 57 (*51*). Finally, since yahs is able to break mis-joined contigs, we performed a last round of purging as described above.

We assessed final assembly quality based on heatmaps of Hi-C contact links and BUSCO scores. To get Hi-C contact heatmaps, we generated hic files using juicer tools v1.1.9 (*52*). We generated heatmaps using HiCExplorer (v3.7.2, hicConvertFormat -outputFormat cool, hicPlotMatrix --log1p (*53*)). We calculated BUSCO scores using insecta_odb10 v5.4.5 database (*54*). We used positions and IDs of single copy and complete BUSCO genes in *B. g. grandii* and *B. r. redtenbacheri* genomes to evaluate synteny between the two genomes.

### Pseudoreference construction

To aid the haplotype phasing of reads from *B. lynceorum* and hybridogen 2 (see Results) we constructed pseudoreferences of *B. atticus* and *B. g. benazzii*. For this we mapped the WGS short reads of *B. atticus* and *B. g. benazzii* to the reference genome of *B. g. grandii* using bwa mem v0.7.17 with default options (*39*). We sorted the alignments using samtools v1.15.1 (*41*) and called variants using bcftools mpileup and bcftools call with the option --multiallelic-caller. We applied the variants to the reference genome of *B. g. grandii* using bcftools consensus (*42*). We applied heterozygous variants as IUPAC characters. For sites with missing data we kept the sequence of *B. g. grandii* unchanged.

All commands, see “tree_genomes.sh” (https://github.com/AlexPopalex/tree/).

### Maternal & paternal haploset trees

#### Phasing RADseq data

We phased RAD reads of all hybrid individuals (see “Species determination”; Supplementary Figure 2) by competitive mapping to the concatenated genomes of the corresponding parental species using bbsplit (bbmap v38.6.3; https://sourceforge.net/projects/bbmap). Specifically, we competitively mapped reads of *B. whitei* and hybridogen 1 to the *B. r. redtenbacheri* and the *B. g. grandii* reference genomes. We mapped reads of hybridogen 2 to the *B. r. redtenbacheri* reference genome and the *B. g. benazzii* pseudoreference. We mapped reads of *B. lynceorum* to the *B. r. redtenbacheri* and the *B. g. grandii* reference genomes as well as the *B. atticus* pseudoreference. We mapped reads of three individuals described as novel hybrids (*23*) to their parental references, i.e. to the *B. g. grandii* reference genome and the *B. atticus* pseudoreference for novel hybrid 1, the *B. g. grandii* reference genome and the *B. g. benazzii* pseudoreference for novel hybrid 2 and the *B. r. redtenbacheri,* the *B. g. grandii* reference genome and the *B. g. benazzii* pseudoreference for novel hybrid 3. We discarded ambiguously mapping reads with the option ambiguous2=toss. We calculated alternative allele depth ratios from the filtered vcf used in species determination (see above) with a custom script. We tested unmapped reads for contamination using kraken v2.1.3 and the standard database (*55*).

To assess the quality of our phasing pipeline, we artificially generated hybrids *in silico* by combining reads from non-hybrid individuals (five randomly chosen individuals of *B. rossius*, *B. g. grandii*, *B. g. benazzii* and *B. atticus*) so as to mimic five hybrid individuals for each relevant combination of genomes representing hybridogen1/*B. whitei*, hybridogen 2 and *B. lynceorum.* These *in silico* hybrids were processed alongside the natural hybrids throughout all downstream analysis (see Suppl. Results).

#### Variant calling

To reconstruct the maternal haploset tree we re-mapped all phased reads from hybrids that competitively mapped to *B. r. redtenbacheri* as well as reads of *B. r. redtenbacheri* and *B. r. rossius* individuals to the *B. r. redtenbacheri* reference genome. To reconstruct the paternal haploset tree we re-mapped all phased reads of hybrids competitively mapping to *B. g. grandii*, *B. g. benazzii* or *B. atticus* as well as reads of *B. grandii* and *B. atticus*, individuals. We sorted alignments using samtools and removed unmapped reads and secondary and supplementary alignments as well as alignments with mapping quality below 20 (*41*). In addition, we filtered the alignments for non-uniquely mapping reads. We called variants using bcftools mpileup and bcftools call (*42*) with the option --multiallelic-caller and including gvcf blocks of homozygous reference calls and missing sites (--gvcf 0). These datasets combine reads from diploid genomes (from non-hybrid individuals) with reads from haploid genomes (phased reads from hybrids). Nevertheless we called all variants in diploid mode to enable identification and removal of false positive heterozygous genotypes of the haplotypes obtained from hybrids. We filtered variants using bcftools (*42*). We removed duplicate records, indels, polyallelic sites (keeping mono- and biallelic sites), sites with variant quality < 20 and sites with a depth > median of per genotype depth + 2* standard deviation of per genotype depth * No. of samples. We replaced genotypes with depth < 4 and any heterozygous genotypes with missing information in order to remove false positive heterozygous sites for the hybrids haplotypes.

#### Tree reconstruction

We applied the maternal and the paternal variants to the reference genomes of *B. r. redtenbacheri* and *B. g. grandii* per sample using bcftools consensus (*42*). We restricted the application to the 18 longest scaffolds for *B. r. redtenbacheri* and the 17 longest scaffolds for *B. g. grandii* representing their karyotypes of 1n=18 and 17, respectively (see “Results: Genome assembly” and (*23*). For sites and genotypes with missing data we changed the reference genome sequences to N. We aligned the 369 maternal and the 372 paternal haplosets per scaffold and removed sites where more than 25% of individuals had missing genotypes from each alignment using trimal (*56*). We concatenated the alignments of all scaffolds using geneious prime 2022.0.1. The resulting maternal and paternal alignments were 2,388,088 bp and 1,925,680 bp long, covering 0.154 and 0.116 % of their assemblies (18 and 17 longest scaffolds), respectively. We reconstructed phylogenetic trees per scaffold and for the concatenated maternal and paternal alignments (to obtain whole-haploset branch length estimates) using iqtree2 with the implemented Model Finder Plus and 1000 bootstrap replicates (*57*). We generated majority rule consensus trees from the per-scaffold trees and visualized them using geneious prime 2022.0.1.

All commands, see “tree_rad.sh” (https://github.com/AlexPopalex/tree/).

### WGS Data

Phasing, variant calling & tree reconstructions:

To phase the PacBio reads of *B. whitei* we combined the reference genomes of *B. r. redtenbacheri* and *B. g. grandii* into one fasta file. We then mapped the reads onto the combined fasta file using minimap2 v2.14 with the option -x map-hifi and allowing for up to 50 secondary alignments (-N 50) (*40*). We filtered the alignment for reads that mapped unambiguously, i.e. either to *B. r. redtenbacheri* or to *B. g. grandii* using samtools v1.15.1 and seqtk v1.4-r122 (*41*) (https://github.com/lh3/seqtk). For phasing of the two lineages of *B. lynceorum* we added the pseudoreference of *B. atticus* to the concatenated reference genomes of *B. r. redtenbacheri* and *B. g. grandii*. We mapped the PacBio reads of *B. lynceorum* as described for *B. whitei* and filtered for reads that mapped unambiguously either to *B. r. redtenbacheri* or to *B. g. grandii* or to the *B. atticus* pseudoreference. To remove potentially contaminating reads of *B. grandii* origin (e.g. reads derived from maternal somatic/follicle cells in the unfertilized eggs; see Results) we also phased the WGS short reads of the hybridogens. Using bbsplit (https://sourceforge.net/projects/bbmap), we competitively mapped reads of hybridogen 1 to the *B. r. redtenbacheri* and *B. g. grandii* reference genomes and of hybridogen 2 to the *B. r. redtenbacheri* reference genome and the *B. g. benazzii* pseudoreference. We discarded ambiguously mapping reads with the option ambiguous2=toss. Variant calling and tree reconstructions were done as described above.

#### Topology testing and divergence estimates

We tested different evolutionary scenarios by comparing the goodness of fit of the best fitting ML tree to the alignment with different constrained topologies. First, we calculated a tree constrained to a topology separating hybridogenesis and parthenogenesis, i.e. where the *rossius* haplome of hybridogen 1 and 2 form a monophyletic clade and the *rossius* haplome of *B. whitei* and both lineages of *B. lynceorum* form a monophyletic clade using iqtree2 with TVM+F+I as the best-fit model. We then tested if the best fitting ML tree was a significantly better fit to the alignment than the constrained tree using RELL approximation with 10,000 RELL replicates and the Shimodaira-Hasegawa test implemented in iqtree2 (*58*). Other tests probing different evolutionary scenarios (see Fig 2 D, E, F) were conducted using the same approach (see Supplementary Results 6).

We calculated an approximate minimum age estimate of the origin of hybridogenesis by dividing the branch length of hybridogen 2 by a germline mutation rate of 2.82 * 10^-9^ (estimate for the most recent common ancestor of arthropods [(*59*)]; *Bacillus* has one generation per year.

To estimate uncorrected pairwise differences we removed all positions containing Ns from the whole haplome alignment and calculated the numbers of pairwise differences using Geneious prime v 2022.0.1.

All commands, see “tree_genomes.sh” (https://github.com/AlexPopalex/tree/)

### Mitochondrial tree

To reconstruct pseudoreferences of the mitochondrial genomes we mapped the genomic reads of *B. r. redtenbacheri* and the hybrid lineages (including two lineages of *B. lynceorum*) to the partial mitochondrial genome of *B. rossius* retrieved from NCBI (GenBank: GU001956.1) (*60*) using bwa and minimap, respectively, and sorted the alignments using samtools (as described above under “Species determination”). We called and filtered variants as described above under “Maternal and paternal haploset trees” (except we did not remove sites exceeding a maximum depth value). Variants were applied to the partial mitochondrial genome for each species, sequences were aligned and sites with any ambiguous bases removed. Finally, a phylogenetic tree was constructed (all as described above under “Maternal and paternal haploset trees”).

### Experimental validation of inferred transitions

We validated the inferred transitions by replicating them experimentally in crosses between different strains. The crosses were originally conducted for purposes unrelated to the present study, but led to serendipitous findings that illustrate the three key transitions following the loss of sex revealed by our phylogenetic approach (see Figure 3A): Transition 2 (“host switch”), replacement of the paternal haplome in hybridogens by that of a new lineage; Transition 3 from hybridogenesis to parthenogenesis, i.e. production of unreduced F1 hybrids; and Transition 4 from diploid to triploid parthenogenesis by incorporation of a third genome by a diploid parthenogen upon fertilization of a small proportion of eggs.

#### Host Switch (Transition 2)

To validate transition 2, we crossed 12 hybridogen 1 females (*B. rossius* x *B. g. grandii*) from our stock populations with five *B. g. benazzii* males. We then genotyped the progeny (n = 282) of these crosses using two microsatellites with fixed differences between the *B. rossius* and *B. grandii* genomes (for both *grandii* and *benazzii* subspecies; see Supplementary Table S2). We used the QIAGEN Multiplex PCR kit (following manufacturer’s instructions) with the following thermocycler conditions: 15m 95°C (initial denaturation); 35 cycles of 30s 94°C (denaturation), 90s 60°C (annealing), 90s 72°C (extension); 10m 72°C (final extension). The PCR products were sent to Microsynth AG for fragment length analysis (using the Mic-G5-RCT dye set). Alleles were called with the open source software OSIRIS developed by NCBI (Available from: https://www.ncbi.nlm.nih.gov/osiris/).

#### Hybridogenesis to parthenogenesis (Transition 3)

For transition 3, we crossed five virgin females of hybridogen 1 with males of *B. rossius* thereby generating F_1_ hybrid females with a “hybridogenetic” and a “sexual” haplome of *B. rossius*. We then backcrossed these females with males of *B. g. grandii* (10 males that were collected in Cassibile in 2020 and 2021, respectively) to restore, in their offspring (F2 hybrids), the original hybrid state between *B. rossius* and *B. grandii* found in the wild hybridogenetic and parthenogenetic lineages. To assess the ability of F_2_ females to reproduce via hybridogenesis, we collected and sequenced a single unfertilized egg for 93 of them. Thus, hybridogenetic reproduction would be indicated by the lack of *B. grandii* genome in the unfertilized embryo. We extracted DNA from the eggs, prepared RADseq libraries as described above and sequenced 150bp paired end reads on one lane of the NovaSeq S4 (yielding 400 M reads in total).

To identify traces of *B. grandii* stemming from the chorion or nurse cells of maternal origin, which would result in erroneous reproductive mode inference, we first compared the depth of reads of *B. rossius* versus *B. g. grandii* origin in sliding windows along both genomes (Figure 3). We concatenated and sorted the 18 longest scaffolds of the *B. r. redtenbacheri* reference genome (note that scaffolds 4 and 9 are split in the reference genome) to be able to represent them in synteny to the *B. g. grandii* reference genome in the coverage comparisons. We then competitively mapped RAD reads of the 93 unfertilised eggs to the synteny-adapted reference genomes using bwa. We only retained primary alignments and properly paired reads using samtools. We calculated sliding window coverage using tinycov v0.4.0 (https://github.com/cmdoret/tinycov) and adjusted window size per scaffold of the *B. r. redtenbacheri* genome to match the corresponding window size in the *B. g. grandii* genome. Following mapping, we excluded 15 eggs based on their low coverage.

Next we inferred the extent of introgression of the “sexual” haplome of *B. rossius* into “hybridogenetic” haplome for the 78 of the 93 F_2_ females with sufficient egg coverage depth. For this, we extracted all reads with primary alignments to the *B. r. redtenbacheri* genome from the above-generated alignments using samtools. We then generated pseudoreferences for the sexual haplome and the haplome of hybridogen 1 using bwa, samtools and bcftools. We competitively mapped the phased *B. rossius* reads to the pseudoreferences and only retained uniquely mapping, properly paired reads using bwa and samtools. We calculated the proportion of reads from the sexual haplome over all *B. rossius* reads using samtools and bash. We found a bimodal distribution of the proportion of reads mapping to the sexual haplome, indicating admixture in some individuals but elimination of the sexual haplome by the F_1_ in others (Figure S7).

All commands, see “tree_crosses.sh” (https://github.com/AlexPopalex/tree/)

#### Transition to triploidy (Transition 4)

To validate transition 4, we mated 24 *B. whitei* females with *B. g. grandii* males. We used a diplex of two microsatellites with fixed differences between the *B. rossius* and *B. grandii* genomes to assess the genotype of 281 hatchlings from the 24 mated *B. whitei* females (see Supplementary Table 2 and previous section for genotyping protocol). Our approach was not stringent as the identification of triploid progeny would only be possible whenever maternal and paternal *B. g. grandii* alleles were different. Yet, we identified one triploid *B. whitei* hatchling that incorporated a second *B. g. grandii* haploset. We confirmed this genetic composition with ddRADseq (as previously described) by comparing the ratio between reads mapping on the *B. r. redtenbacheri* genome and those mapping on the *B. g. grandii* genome (i.e., twice as many). Such transition from parthenogenetic *B. whitei* to triploidy by incorporation of an additional *B. g. grandii* haploset has already been described (using allozyme polymorphism) and shown to be stable for several generations (*30*). Since *B. whitei* and *B. g. grandii* males are naturally co-occurring in the wild, such interspecific crosses might also happen in natural populations. Thus, we screened all hybrid females collected from the field for signatures of triploidy (n = 155, ddRAD-seq), but we did not identify any triploids. It is currently unclear whether this stems from transitions being too rare to be detected in our set of hybrids, interspecific matings not occurring in natural populations, or from a high mortality of triploids.

**Supplementary Table S2:**
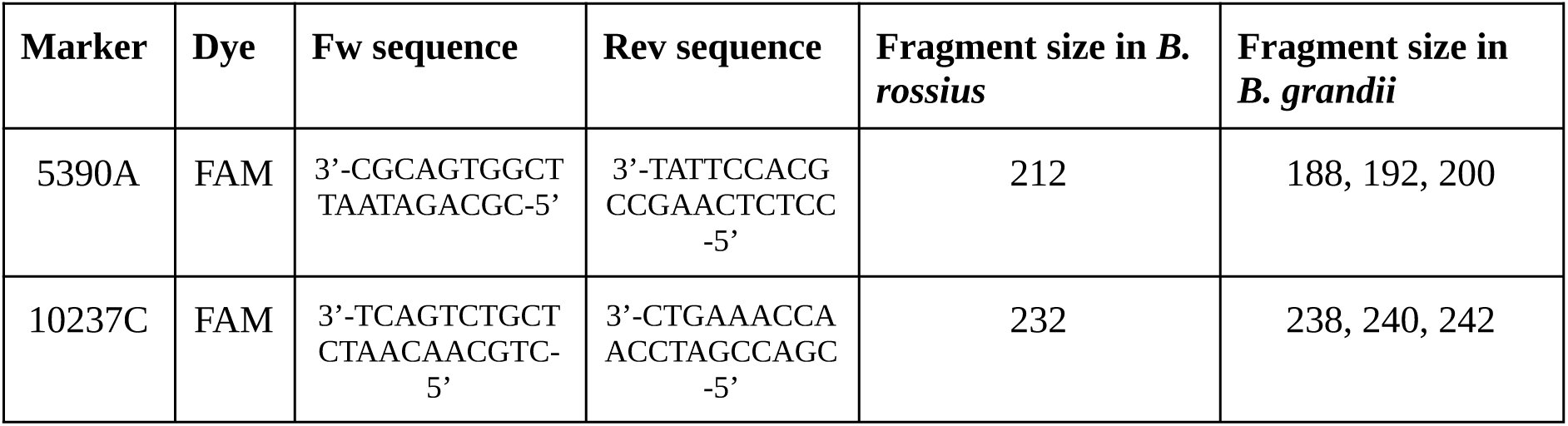
Sequences of primers used to discriminate *B. rossius* and *B. grandii* alleles.

## Supplementary Results

### Suppl. Results 1: Species Determination

*Bacillus* species and lineages are morphologically indistinguishable, therefore we identified individuals by comparing their genotypes to the genotypes of previously identified individuals (*23*) (see Figure S1). This approach allowed for the distinction of nine lineages, with hybridogen1 and *B. whitei* merged into a single lineage as a consequence of their similar genome composition: *B. rossius*: 184 inds. RADseq (+1 ind. whole genome), *B. g. grandii*: 74 (+1), *B. g. benazzii*: 53 (+1), *B. g. maretimi*: 8, *B. atticus*: 13 (+1), hybridogen1/*B. whitei*: 102 (+2), *B. lynceorum*: 29 (+1), hybridogen2: 38 (+1), novel hybrids: 3 inds.). Hybridogen1 and *B. whitei* were clearly split in the phylogenetic tree and identified using previous information on reproductive mode of related individuals (*23*): hybridogen1: 47 (+1), *B. whitei*: 55 (+1). The different hybrid lineages were female-only or characterized by highly female-biased sex ratios (inferred from RADseq data): hybridogen 1/*B. whitei:* 2 males, 101 females, 1 NA; hybridogen 2: 12 males, 25 females, 1 NA (note that read phasing later revealed that 20 of these are novel F_1_ hybrids between *B. r. redtenbacheri* females and *B. g. benazzii* see below). Finally, the genomic composition of three novel hybrids (a, b, c; black points) described in (*23*) was confirmed.

**Figure S1:**
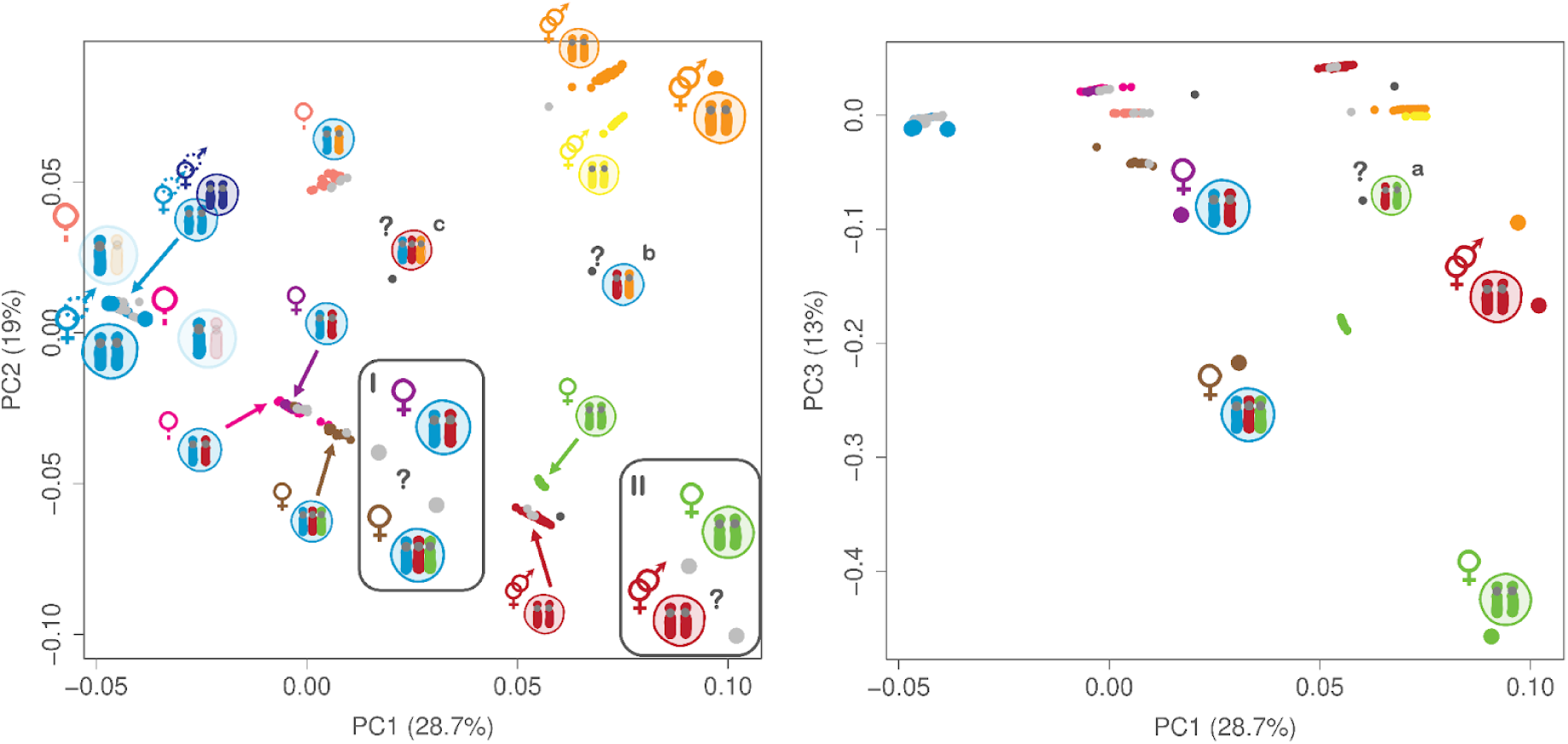
Genetic species determination of 512 *Bacillus* individuals using PCA. Small points represent 504 genotypes inferred from RADseq data. Colored points represent samples with known species identity (*23*). The genomic composition and reproductive strategy of clusters is indicated as small symbols; see Figure 1 C for species names and symbol legend. Grey points represent samples that were added in this study. Large coloured points indicate eight genotypes inferred from whole genome sequencing data. PC1 reflects genetic distance between the subspecies of *B. rossius* and the subspecies of *B. grandii*, PC2 reflects the distance among subspecies of *B. grandii* and PC3 reflects the distance between *B. atticus* and *B. grandii*. PC3 was used to differentiate between hybridogen1/*B. whitei* and the triploid *B. lynceorum* (right plot). Note that 15 artificial *in silico* hybrids used to assess phasing quality (see methods) were removed from the plot. Large grey points with question marks indicate that species assignment was ambiguous between two species (*B. g. grandii* vs. *B. atticus* and *B. lynceorum* vs. *B. whitei*) in the plot of PC1 vs PC2 (left) but could be resolved in the plot of PC1 vs PC3 (right).

### Suppl. Results 2: Genome Assembly

To enable the phasing of data from hybrid lineages, we assembled a high quality genome of *B. g. grandii* (BUSCO: C:99.0%[S98.5%,D:0.5%],F:0.7%, M:0.3%, n:1367; see Methods). The genome is 1.7 Gb (haploid genome size) with 96.64% anchored to 17 super-scaffolds, ranging in size from 27 to 441 Mb (see Figure S2).

**Figure S2:**
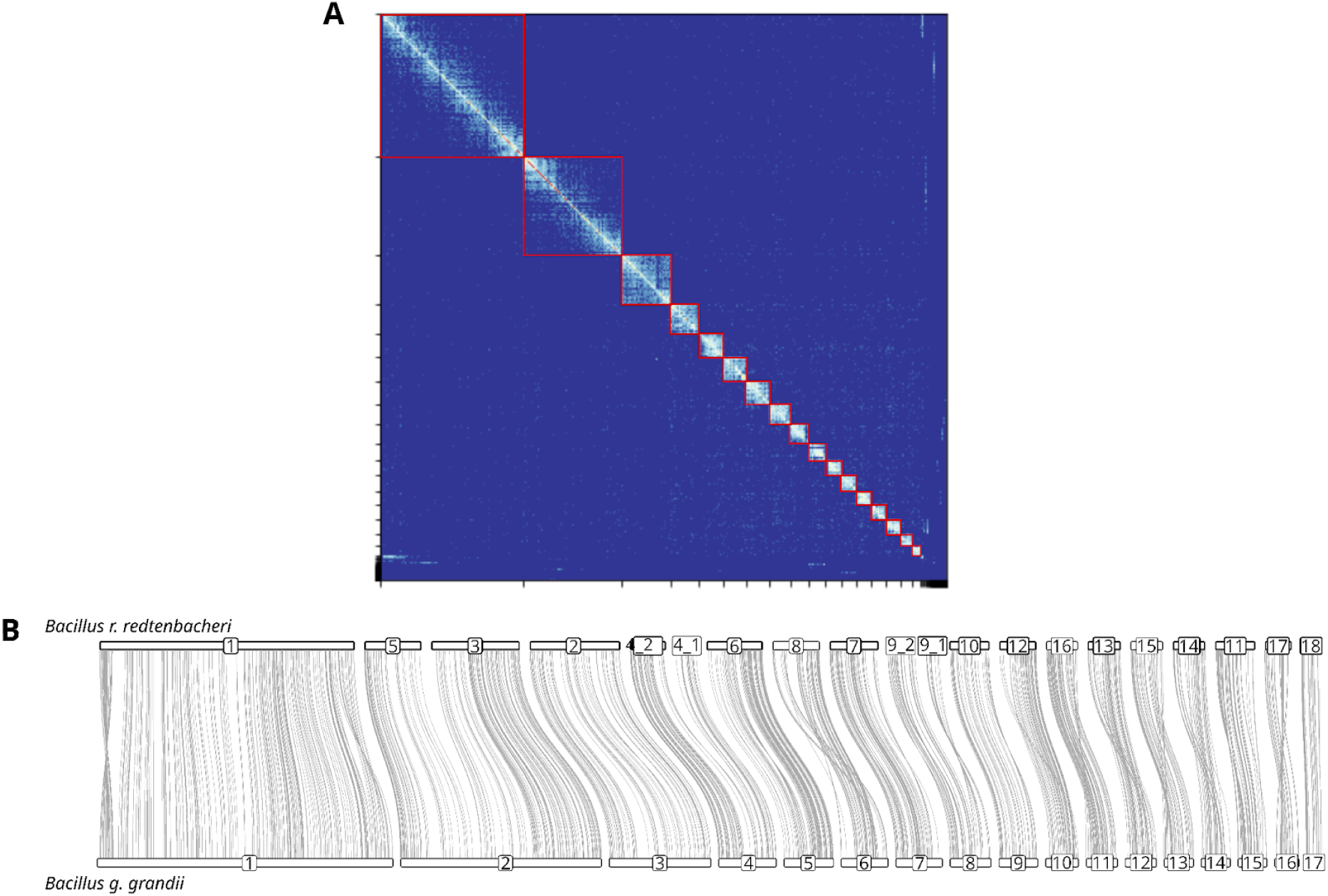
**A.** Hi-C contact map for the *B. g. grandii* assembly. The dark blue to red scale denotes contact intensity and red squares highlight super-scaffolds. **B.** Syntenic relationships between *B. r. redtenbacheri* and *B. g. grandii* genomes. Links between genomes correspond to orthologous BUSCO genes.

### Suppl. Results 3: Pseudoreference Construction & Phasing

Using whole genome sequencing reads of *B. atticus* and *B. g. benazzii* and the assembled genome of *B. g. grandii* as a template we constructed pseudoreferences in order to aid phasing of *B. lynceorum* and hybridogen2, respectively. For the phasing of hybridogen1 and *B. whitei* we used the reference genome assemblies of *B. g. grandii* and *B. r. redtenbacheri*. Pseudoreferences were 73.9% (*B. atticus*) and 76.3% (*B. g. benazzii*) complete, i.e. 1,272,962,102 and 1,314,166,200 variants were applied to the 1,721,945,181 bp *B. g. grandii* reference genome, respectively. For sites with missing homologous sequence data (26.1% and 23.7%, respectively), the sequence of *B. g. grandii* was kept unchanged. We phased the whole genome and RADseq reads from hybrid individuals by competitive mapping to the reference or pseudoreference genomes of their maternal and paternal ancestors. RADseq reads of hybridogen1/*B. whitei* mapped to 50.46% and 46.08% (median values; with 4.09 and 4.0 standard deviations) unambiguously to *B. r. redtenbacheri* (maternal ancestor) and *B. g. grandii* (paternal ancestor), respectively. RADseq reads of hybridogen2 mapped to 52.99% and 40.93% (median values; with 1.65 and 0.99 standard deviations) unambiguously to *B. r. redtenbacheri* (maternal ancestor) and the *B. g. benazzii* pseudoreference (paternal ancestor), respectively. The lower percentage of reads mapping to *B. g. benazzii* likely reflects the incompleteness of the *B. g. benazzii* pseudoreference. RADseq reads of *B. lynceorum* mapped to 34.75%, 28.91% and 19.5% (median values; with 5.43, 2.72 and 1.94 standard deviations) unambiguously to *B. r. redtenbacheri* (maternal ancestor), *B. g. grandii* and the *B. atticus* pseudoreference (paternal ancestors), respectively. The low percentage of reads mapping to *B. atticus* likely reflects the incompleteness of its pseudoreference. We also phased the reads of three novel hybrid individuals discovered previously (*23*) (see Figure S1; marked as a,b,c) by mapping them competitively against the genomes/pseudoreference of their respective ancestral species. Reads of novel hybrid a mapped to 48.66% unambiguously to *B. g. grandii* and to 30.33% to *B. atticus*, reads of novel hybrid b mapped to 40.84% to *B. g. grandii*, and to 35.32% to *B. g. benazzii* and reads of novel hybrid c mapped to 37.94% to *B. r. redtenbacheri*, to 25.05% to *B. g. grandii*, and to 20.07% to *B. g. benazzii*.

There were three (out of 102) noticeable outliers of hybridogen1/*B. whitei* (see Figure S3). Two individuals were most likely triploids, with one haploset being derived from *B. r. redtenbacheri* and two from *B. g. grandii*. This was indicated by mapping percentages strongly skewed in favor of *B. g. grandii* (30.16% to 63.77% and 34.9% to 60.53% of unambiguously mapping reads) and heterozygous genotypes with an excess alternative allele depth ratio for the paternal ancestor *B. g. grandii* (0.62 and 0.59, respectively). The third outlier showed low mapping percentages (26.81% and 20.09%) for both ancestral genomes, with 52.04% non mapping reads, and an unbiased alternative allele depth of 0.49. We found a high percentage (24.1%) of the reads being of bacterial origin and derived from contamination (see Methods).

**Figure S3:**
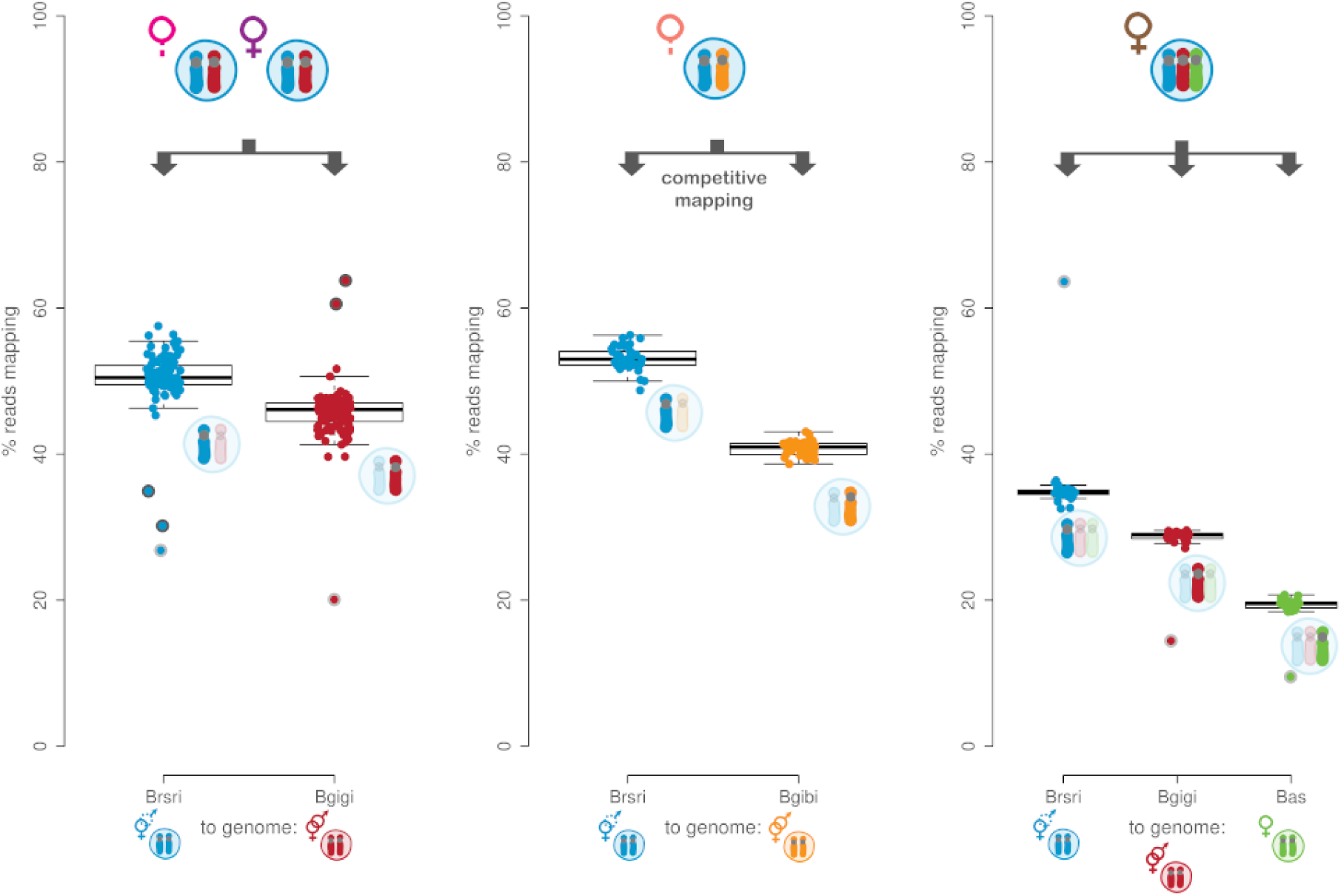
Phasing of RADseq reads via competitive mapping. Reads from individuals of the different hybrid species (symbolised on top; see Figure 1C) were mapped competitively (indicated as arrows) to the reference genomes (or pseudoreferences) of their parental species. This enabled the separation of reads derived from the different haplomes (highlighted below the whiskers). A) RADseq reads of hybridogen1/*B. whitei* were mapped competitively to the reference genomes of *B. r. redtenbacheri* (Brsri) and *B. g. grandii* (Bgigi). Each individual is represented as two points, with a mapping percentage to Brsri (blue point) and a mapping percentage to Bgigi (red point). Two individuals showed mapping percentages indicative of triploidy with one haploset of Brsri and two haplosets of Bgigi (dark grey circles). One individual showed mapping percentages that were uniformly reduced for both reference genomes (likely due to contamination; light grey circle). B) RADseq reads of hybridogen 2 were mapped competitively to the reference genomes of *B. r. redtenbacheri* (Brsri) and the pseudoreference of *B. g. benazzii* (Bgibi). Each individual is represented as two points, with a mapping percentage to Brsri (blue point) and a mapping percentage to Bgibi (orange point). C) RADseq reads of *B. lynceorum* (symbolised; see Figure 1C) were mapped competitively to the reference genomes of *B. r. redtenbacheri* (Brsri), *B. g. grandii* (Bgigi) and the pseudoreference of *B. atticus* (Bas). Each individual is represented as three points, with a mapping percentage to Brsri (blue point), a mapping percentage to Bgigi (red point) and a mapping percentage to Bas (green point). The outlier (light grey circle) is likely the result of cross contamination (as indicated by the position of the individual in the maternal tree).

There was one apparent outlier of *B. lynceorum* with mapping percentages strongly skewed in favor of *B. r. redtenbacheri* (63.59% to 14.43% to 9.49%). The position of the outlier in the phylogenetic tree suggests that the skewed mapping percentages are derived from contamination. The four outliers were not included in any further analyses.

### Suppl. Results 4: Maternal Tree

We calculated a haplome tree based on SNPs from maternal RADseq reads that were mapped to the reference genome of *B. r. redtenbacheri* (maternal ancestor of all hybrids). The tree comprised 369 tips, 184 representing individuals of different *B. rossius* lineages from Sicily and the Italian (and French) mainland, and 185 representing the phased maternal haplosets (hereafter referred to as *rossius* haplosets) of hybrid individuals (see Figure S4).

Additionally, the tree comprised 15 “artificial *in silico* hybrids” (see methods), which we introduced in order to test whether incomplete phasing could influence the position of hybrids in the phylogenetic tree. If phasing was complete, we expect their *rossius* haplosets to cluster next to the five *B. rossius* individuals that we used as a source. This was indeed the case as all 15 *rossius* haplosets clustered next to their *B. rossius* source individuals.

The *rossius* haplosets of all *B. whitei* (55 individuals), *B. lynceorum* (28), hybridogen 1 (47) as well as 18 out of 38 individuals of hybridogen 2 formed a single clade (Figure S4). Surprisingly, the remaining 20 individuals (12 males, 7 females, 1 NA) identified as hybridogen 2 (based on unphased genome data) were interspersed among *B. r. redtenbacheri* females from Scopello, indicating that they are in fact newly-formed hybrids between *B. r. redtenbacheri* and *B. g. benazzii*.

The majority of the remaining *B. r. redtenbacheri* formed a separate clade with the populations from the italian mainland, Forgiano (15 inds.) and Ravenna (10 inds.) separating from the Sicilian populations, Cefalu (11 inds.) and Patti (25 inds.). The individual of *B. lynceorum* with skewed mapping percentages (see above) clustered with the individuals from Patti, presumably indicating cross contamination between one *B. lynceorum* individual and one *B. r. redtenbacheri* individual from Patti. Two *B. r. redtenbacheri* individuals from Gorghi Tondi and one from C. di Mazara (both Sicily) formed a sister group to all other *B. r. redtenbacheri* and the haplosets of the hybrids. Individuals from a population sampled in a region of potential overlap in distribution between *B. r. redtenbacheri* and *B. r. rossius* near the Vesuve mountain formed one clade (25 inds.; see Figure 1D; (*21*)). Finally, individuals from an asexual *B. r. rossius* population from Pise (19 inds.), 3 female individuals from Grosseto as well as the sexual *B. r. rossius* population from La Ciotat (France; 18 inds.) formed separate clades.

**Figure S4:**
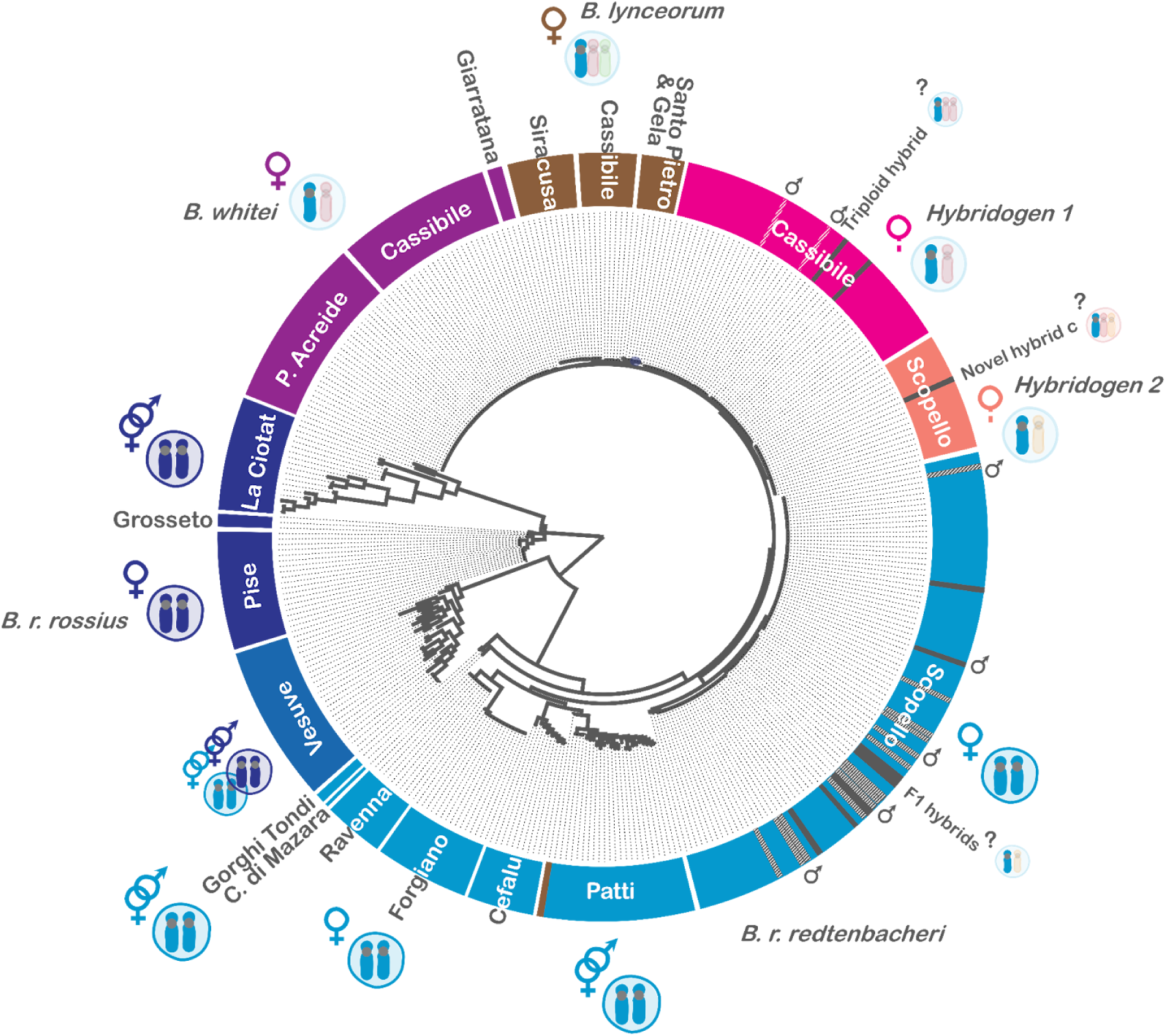
Maternal tree displaying phylogenetic relationships of 369 individuals and maternal haplosets (184 inds. of *B. rossius* and 185 *rossius* haplosets of hybrids). (Sub)species are colored according to Figure 1 C and their populations highlighted around the tree. Genomic composition and reproductive modes of (sub)species are symbolized (with the *rossius* haplosets highlighted). Grey bars indicate two triploid females of hybridogen 1, novel hybrid c and putative F1 hybrids from Scopello. White striped bars indicate two males among hybridogen 1 and putative F1 hybrid males from Scopello. One individual of *B. lynceorum* in the Patti population likely derives from cross contamination (see Results).

### Suppl. Results 5: Paternal Tree

We calculated a haploset tree based on SNPs from paternal RADseq reads that were mapped to the reference genome of *B. g. grandii*. The tree comprised 372 tips, 148 representing individuals of different *B. grandii* subspecies and *B. atticus* from Sicily, and 224 representing the phased paternal haplosets derived from *B. g. grandii* (hereafter referred to as *grandii* haploset), *B. g. benazzii* (*benazzii* haploset) and *B. atticus* (*atticus* haploset) of hybrids (see Figure S5).

The 20 paternal haplosets of the artificial *in silico* hybrids clustered next to their *B. grandii* and *B. atticus* “source” individuals, respectively, indicating that our phasing pipeline accurately separated reads deriving from different haplomes.

All *grandii* haplosets, i.e. those of *B. whitei* (55 inds.), *B. lynceorum* (29 inds.), hybridogen1 (47 inds.) and *B. g. grandii* (74 inds.) formed one monophyletic clade (Figure S5). This clade also comprised the *B. g. grandii* haplosets of the novel hybrids a, b and c as well as the two triploid hybrids. Individuals of *B. g. grandii* carrying a mitochondrial genome of *B. r. redtenbacheri* origin (likely derived from androgenesis; 15 inds.) formed multiple small clades or were placed isolated among *B. g. grandii* indicating the absence of mating preferences between cytoplasmic hybrids.

The *atticus* haplosets of *B. lynceorum* were paraphyletic, indicating that the two lineages of *B. lynceorum* originated independently by hybridisation with different lineages of *B. atticus.* Individuals from Santo Pietro and Gela (*B. lynceorum* 2) formed a monophyletic clade together with the individuals of *B. atticus* (13 inds.; and the *atticus* haploset of novel hybrid a), while individuals from Siracusa and Cassibile (*B. lynceorum* 1) formed another monophyletic group.

The *benazzii* haplosets of hybridogen 2 (female only; 18 inds.) and the putative F1 hybrids (males and females; 20 inds.; see above) formed a clade with their ancestral subspecies *B. g. benazzii* (53 inds.). This clade also comprised the *benazzii* haplosets of the novel hybrids b and c. Note that *B. grandii* was paraphyletic with *B. g. grandii* forming a sister group with *B. atticus* consistent with topologies based on partial COII sequences (*21*, *23*).

**Figure S5:**
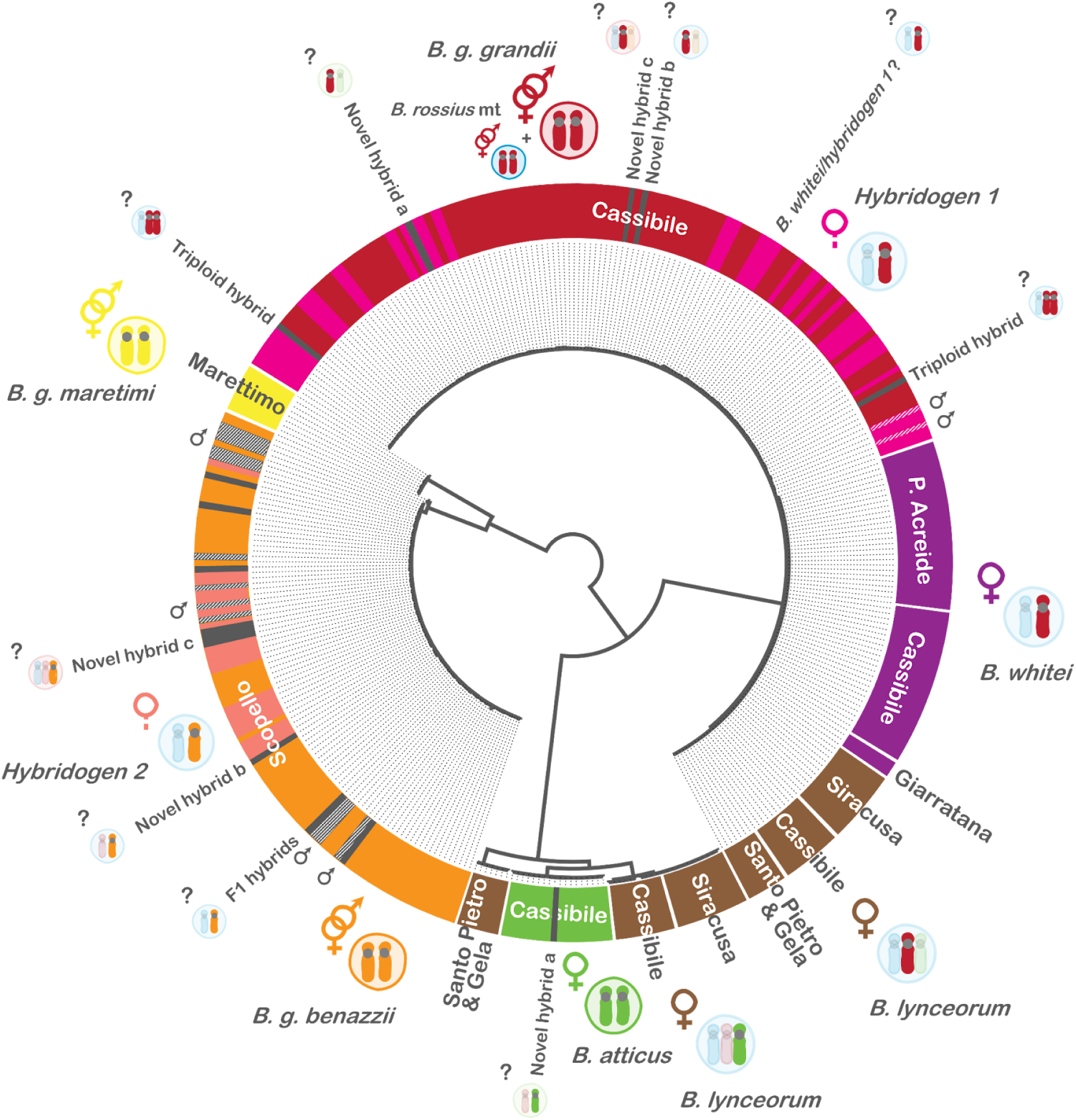
Paternal tree displaying the phylogenetic relationships of 372 individuals and paternal haplosets (148 inds. of *B. grandii* and *B. atticus*, and 224 *grandii*, *benazzii* and *atticus* haplosets of hybrids). (Sub)species are colored according to Figure 1C and their geographical locations are highlighted around the tree. Genomic composition and reproductive modes of (sub)species are symbolized with the paternal haplosets highlighted. Grey bars indicate two triploid females of hybridogen 1, the novel hybrids a, b, c and putative F1 hybrids from Scopello. White striped bars indicate two males among hybridogen 1 and putative F1 hybrid males from Scopello.

### Suppl. Results 6: Whole genome & mitochondrial tree

The RADseq-based maternal tree did not allow us to confidently estimate the order of transitions between alternative reproductive strategies. To increase the number of phylogenetically informative sites we sequenced the genome of *B. atticus*, *B. g. benazzii*, *B. whitei* and the two lineages of *B. lynceorum*, and phased the reads using competitive mapping. We also sequenced the *B. r. redtenbacheri* haplomes of both hybridogens using unfertilised eggs (see Figure 1B and Methods).

Competitive mapping for *B. whitei* unambiguously attributed 47.7% of reads to *B. r. redtenbacheri* and 50% to *B. g. grandii* (2,572,341 and 2,696,990 of 5,391,303 mapped reads). Competitive mapping for the *B. lynceorum* lineage from Siracusa (*B. lynceorum* 1) unambiguously attributed 30% of reads to *B. r. redtenbacheri*, 23% to *B. g. grandii* and 8.6% to *B. atticus*. (496,698 and 382,032 and 142,107 of 1,657,689 mapped reads). Competitive mapping for the *B. lynceorum* lineage from Gela (*B. lynceorum* 2) unambiguously attributed 30.4% of reads to *B. r. redtenbacheri*, 21.93% to *B. g. grandii* and 8.52% to *B. atticus*. (1,768,358 and 1,275,673 and 495,473 of 5,817,866 mapped reads).

We also phased the reads of the hybridogens in order to remove DNA sequences potentially derived from introgression (leaky hybridogenesis; but see (*23*)) of *B. grandii* or from retained somatic cells like follicle cells. Indeed, for hybridogen 1 we found 3.3% of read pairs unambiguously attributed to *B. g. grandii* and for hybridogen 2 0.4% of read pairs attributed to *B. g. benazzii* and removed them from phylogenetic tree reconstruction.

We calculated a tree based on SNPs from whole genome sequencing reads of the parental (sub)species *B. r. redtenbacheri*, *B. g. grandii*, *B. g. benazzii* and *B. atticus*, of the phased maternal and paternal haplomes of *B. whitei* (*rossius* haplome and *grandii* haplome from here) and the two lineages of *B. lynceorum* (with the *atticus* haplome in addition), and of the maternal haplosets of the two hybridogens that were mapped to the reference genome of *B. r. redtenbacheri*. We obtained an approximate minimum age estimate of the origin of hybridogenesis of ∼8,000 years (8,156 years to be precise, indicative of a post ice age origin) by dividing the branch length of hybridogen 2 (2.3 * 10^-5^) by a germline mutation rate of 2.82 * 10^-9^ (estimate for the most recent common ancestor of arthropods (*59*); *Bacillus* has one generation per year).

The best fitting ML topology was a significantly better fit to the whole genome sequence alignment than a tree fixed to separate hybridogenesis and parthenogenesis (p < 0.001; SH test). The best fitting ML tree was also a significantly better fit to the whole genome sequence alignment than a tree fixed to separate *B. whitei* from *B. lynceorum* (for the *rossius* haplome of *B. lynceorum* p < 0.001; for the *grandii* haplome of *B. lynceorum* p < 0.001; SH test).

The mitochondrial phylogenetic tree was star-shaped with a single nucleotide polymorphism separating *B. r. redtenbacheri* and hybridogen 2 from the hybridogen 1 and the parthenogenetic lineages (Figure S6).

**Figure S6:**
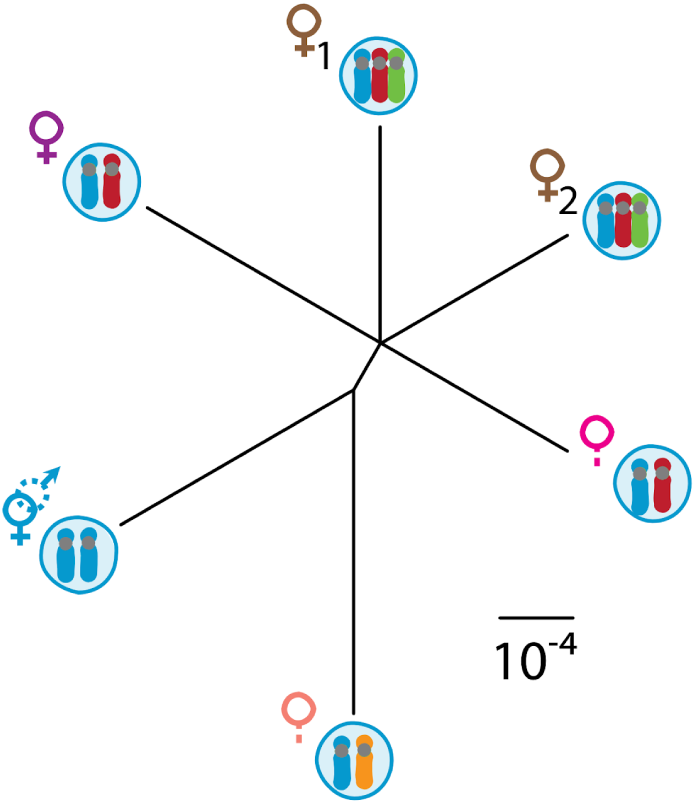
Phylogeny of the whole mitochondrial sequence of the five hybrid lineages and *B. r. redtenbacheri*. *B. r. redtenbacheri* and hybridogen 2 are separated from the other lineages by one substitution.

### Suppl. Results 7: Empirical evidence for the inferred transitions

#### Host switch (Transition 2)

All 228 genotyped female offspring retained their *B. rossius* allele but replaced their *B. g. grandii* alleles with new alleles, which undoubtedly came from their *B. g. benazzii* fathers (for which we did not have genotypes). Two alleles were present at the X-linked locus 5390A. All offspring from one father had one allele (192) while all offspring from the other 4 had a different allele (189).

#### Transition from hybridogenesis to parthenogenesis (Transition 3)

Out of the 93 unfertilized eggs sequenced, 78 had enough coverage to allow us to assess the relative coverage of the two haplomes along the genome. We observed three categories of eggs: 1) “Hybridogenesis” (33 eggs, 42.3%): coverage of the *B. grandii* haplome nearing 0 (sometimes slightly above 0 most likely due to the inclusion of maternal somatic cells in the egg but much lower than *B. rossius*), indicative of normal hybridogenesis with full elimination of the *grandii* haplome. 2) “Aneuploid” (12 eggs, 15.4%): whenever evidence for recombination was observed (i.e. relative coverage of the two haplomes changing abruptly in the middle of a chromosome; see for example chromosome 2 on the middle panel of Figure 3C), the two haplomes had inconsistent coverage along the genome, with some chromosomes or chromosome sections present in more or fewer than two copies. Interestingly, this suggests that hybridogenesis in *Bacillus* is an “all or nothing” process: genome elimination is either complete or absent, but intermediate states such as chromosome-specific elimination of the *grandii* haplome, which would be indicative of a per-chromosome mechanism of hybridogenesis, was never observed. 3) “Parthenogenesis” (33 eggs, 42.3%): equal coverage of both *B. rossius* and *B. grandii* haplomes throughout the genome, indicative of the failure to eliminate the *B. grandii* haplome.

We isolated reads of *B. rossius* origin in order to infer the extent of introgression of the “sexual” haplome of *B. rossius* into the “hybridogenetic” haplome in the eggs, which would be caused by segregation and recombination between the two haplomes in the F_1_ hybrids. Twenty-three eggs showed a proportion of introgressed “sexual” haplome ranging between 0.2 and 0.4 indicating introgression (Figure S7). Out of them, seven matched the “Aneuploid” category (see above and Figure 3C) and 16 matched the “Parthenogenesis” category. The absence of eggs matching the “Hybridogenesis” category suggests that elimination of the *B. grandii* genome always failed when the *rossius* haplome is introgressed, resulting either in aneuploidy or a “rescue” of the eggs via a transition to parthenogenesis.

**Figure S7:**
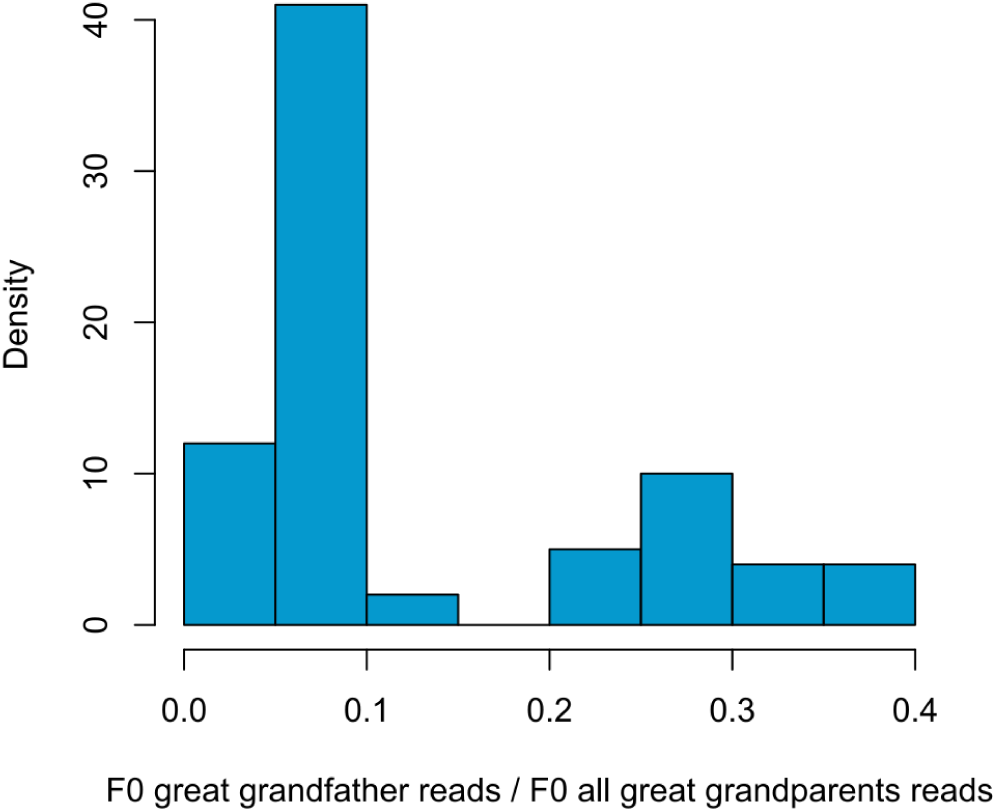
Histogram indicating two categories of unfertilized eggs regarding the proportion of reads derived from the F0 great grandfathers. 55 eggs were 0-0.15 and 23 eggs were 0.2-0.4.

## References

1. R. C. Vrijenhoek, Animal Clones and Diversity. Bioscience 48, 617–628 (1998).

2. B. B. Normark, The evolution of alternative genetic systems in insects. Annu. Rev. Entomol. 48, 397–423 (2003).

3. G. Lavanchy, T. Schwander, Hybridogenesis. Curr. Biol. 29, 539 (2019).

4. J. Lehtonen, M. D. Jennions, H. Kokko, The many costs of sex. Trends Ecol. Evol. 27, 172–178 (2012).

5. S. Meirmans, P. G. Meirmans, L. R. Kirkendall, The costs of sex: facing real-world complexities. Q. Rev. Biol. 87, 19–40 (2012).

6. M. Neiman, T. F. Sharbel, T. Schwander, Genetic causes of transitions from sexual reproduction to asexuality in plants and animals. J. Evol. Biol. 27, 1346–1359 (2014).

7. J.-C. Simon, F. Delmotte, C. Rispe, T. Crease, Phylogenetic relationships between parthenogens and their sexual relatives: the possible routes to parthenogenesis in animals. Biol. J. Linn. Soc. Lond. 79, 151–163 (2003).

8. J. Engelstädter, Constraints on the evolution of asexual reproduction. Bioessays 30, 1138–1150 (2008).

9. M. Maraun, M. Heethoff, K. Schneider, S. Scheu, G. Weigmann, J. Cianciolo, R. H. Thomas, R. A. Norton, Molecular phylogeny of oribatid mites (Oribatida, Acari): evidence for multiple radiations of parthenogenetic lineages. Exp. Appl. Acarol. 33, 183–201 (2004).

10. M. Liegeois, M. Sartori, T. Schwander, Extremely widespread parthenogenesis and a trade-off between alternative forms of reproduction in mayflies (Ephemeroptera). J. Hered. 112, 45–57 (2021).

11. T. Schwander, L. Soldini, R. P. Boisseau, V. Mérel, W. S. Toubiana, G. Lavanchy, On the repeated evolution of parthenogenesis in stick insects, EcoEvoRxiv (2025). https://ecoevorxiv.org/repository/view/8766/.

12. T. Schwander, B. J. Crespi, Multiple direct transitions from sexual reproduction to apomictic parthenogenesis in Timema stick insects. Evolution 63, 84–103 (2009).

13. L. Ross, N. B. Hardy, A. Okusu, B. B. Normark, Large population size predicts the distribution of asexuality in scale insects: Distribution of asexuality in scale insects. Evolution 67, 196–206 (2013).

14. A. J. Barley, A. Nieto-Montes de Oca, N. L. Manríquez-Morán, R. C. Thomson, The evolutionary network of whiptail lizards reveals predictable outcomes of hybridization. Science 377, 773–777 (2022).

15. L. Ross, A. J. Mongue, C. N. Hodson, T. Schwander, Asymmetric inheritance: The diversity and Evolution of non-Mendelian reproductive strategies. Annu. Rev. Ecol. Evol. Syst. 53, 1–23 (2022).

16. J. D. Wetherington, K. E. Kotora, R. C. Vrijenhoek, A test of the spontaneous heterosis hypothesis for unisexual vertebrates. Evolution 41, 721–731 (1987).

17. A. A. Lutes, D. P. Baumann, W. B. Neaves, P. Baumann, Laboratory synthesis of an independently reproducing vertebrate species. Proc. Natl. Acad. Sci. U. S. A. 108, 9910–9915 (2011).

18. D. Hojsgaard, J. Greilhuber, M. Pellino, O. Paun, T. F. Sharbel, E. Hörandl, Emergence of apospory and bypass of meiosis via apomixis after sexual hybridisation and polyploidisation. New Phytol. 204, 1000–1012 (2014).

19. C. Larose, G. Lavanchy, S. Freitas, D. J. Parker, T. Schwander, Facultative parthenogenesis: a transient state in transitions between sex and obligate asexuality in stick insects? Peer Community J. 3 (2023).

20. V. Scali, A “ghost” phasmid appearance: the male *Bacillus atticus*(Insecta: Phasmatodea). Ital. J. Zool. (Modena) 80, 227–232 (2013).

21. B. Mantovani, M. Passamonti, V. Scali, The mitochondrial cytochrome oxidase II gene in Bacillus stick insects: ancestry of hybrids, androgenesis, and phylogenetic relationships. Mol. Phylogenet. Evol. 19, 157–163 (2001).

22. B. Mantovani, V. Scali, Hybridogenesis and androgenesis in the stick-insect Bacillus rossius-grandii benazzii (Insecta, Phasmatodea). Evolution 46, 783–796 (1992).

23. G. Lavanchy, A. Brandt, M. Bastardot, Z. Dumas, M. Labédan, M. Massy, W. Toubiana, P. Tran Van, A. Luchetti, V. Scali, B. Mantovani, T. Schwander, Evolution of alternative reproductive systems in Bacillus stick insects. Evolution 78, 1109–1120 (2024).

24. F. Tinti, B. Mantovani, V. Scali, Reproductive features of homospecific hybridogenetically-derived stick insects suggest how unisexuals can evolve. J. Evol. Biol. 8, 81–92 (1995).

25. H. Munehara, M. Horita, M. R. Kimura-Kawaguchi, A. Yamazaki, Origins of two hemiclonal hybrids among three Hexagrammos species (Teleostei: Hexagrammidae): genetic diversification through host switching. Ecol. Evol. 6, 7126–7140 (2016).

26. O. Marescalchi, V. Scali, New DAPI and FISH findings on egg maturation processes in related hybridogenetic and parthenogenetic Bacillus hybrids (Insecta, Phasmatodea). Mol. Reprod. Dev. 60, 270–276 (2001).

27. J. P. Bogart, K. Bi, J. Fu, D. W. A. Noble, J. Niedzwiecki, Unisexual salamanders (genus Ambystoma) present a new reproductive mode for eukaryotes. Genome 50, 119–136 (2007).

28. S.-S. Choi, A. Mc Cartney, D. Park, H. Roberts, T. Brav-Cubitt, C. Mitchell, T. R. Buckley, Multiple hybridization events and repeated evolution of homoeologue expression bias in parthenogenetic, polyploid New Zealand stick insects. Mol. Ecol. 34, e17422 (2025).

29. M. Neiman, D. Paczesniak, D. M. Soper, A. T. Baldwin, G. Hehman, Wide variation in ploidy level and genome size in a New Zealand freshwater snail with coexisting sexual and asexual lineages: Ploidy, sex, and gender in a freshwater snail. Evolution 65, 3202–3216 (2011).

30. F. Tinti, V. Scali, Androgenetics and triploids from an interacting parthenogenetic hybrid and its ancestors in stick insects. Evolution 50, 1251–1258 (1996).

31. L. Comai, The advantages and disadvantages of being polyploid. Nat. Rev. Genet. 6, 836–846 (2005).

32. O. Marescalchi, L. P. Pijnacker, V. Scali, Cytology of Parthenogenesis in Bacillus whitei and Bacillus lynceorum (Insecta Phasmatodea). Invertebr. Reprod. Dev. 20, 75–81 (1991).

33. C. Moritz, W. M. Brown, L. D. Densmore, S. Vyas, S. Donnellan, M. Adams, P. Baverstock, “Genetic diversity and the dynamics of hybrid parthenogenesis in Cnemidophorus (Teiidae) and Heteronotia(Gekkonidae)” in Evolution and Ecology of Unisexual Vertebrates, R. M. Dawley, J. P. Bogart, Eds. (New York State Museum, Albany, NY, 1989), pp. 87–112.

34. B. Mantovani, F. Tinti, M. Barilani, V. Scali, Current reproductive isolation between ancestors of natural hybrids in Bacillus stick insects (Insecta: Phasmatodea). Heredity (Edinb.) 77, 261–268 (1996).

35. M. Stöck, K. P. Lampert, D. Möller, I. Schlupp, M. Schartl, Monophyletic origin of multiple clonal lineages in an asexual fish (*Poecilia formosa*): SINGLE ORIGIN OF MULTIPLE POECILIA CLONES. Mol. Ecol. 19, 5204–5215 (2010).

36. P. Lamelza, J. M. Young, L. M. Noble, L. Caro, A. Isakharov, M. Palanisamy, M. V. Rockman, H. S. Malik, M. Ailion, Hybridization promotes asexual reproduction in Caenorhabditis nematodes. PLoS Genet. 15, e1008520 (2019).

37. L. Choleva, K. Janko, K. De Gelas, J. Bohlen, V. Šlechtová, M. Rábová, P. Ráb, Synthesis of clonality and polyploidy in vertebrate animals by hybridization between two sexual species. Evolution 66, 2191–2203 (2012).

38. M. Martin, Cutadapt removes adapter sequences from high-throughput sequencing reads. EMBnet J. 17, 10 (2011).

39. H. Li, R. Durbin, Fast and accurate long-read alignment with Burrows-Wheeler transform. Bioinformatics 26, 589–595 (2010).

40. H. Li, Minimap2: pairwise alignment for nucleotide sequences. Bioinformatics 34, 3094–3100 (2018).

41. H. Li, B. Handsaker, A. Wysoker, T. Fennell, J. Ruan, N. Homer, G. Marth, G. Abecasis, R. Durbin, 1000 Genome Project Data Processing Subgroup. 2009. The sequence alignment/map format and samtools. Bioinformatics 25, 2078–2079 (2009).

42. H. Li, A statistical framework for SNP calling, mutation discovery, association mapping and population genetical parameter estimation from sequencing data. Bioinformatics 27, 2987–2993 (2011).

43. S. Purcell, B. Neale, K. Todd-Brown, L. Thomas, M. A. R. Ferreira, D. Bender, J. Maller, P. Sklar, P. I. W. de Bakker, M. J. Daly, P. C. Sham, PLINK: a tool set for whole-genome association and population-based linkage analyses. Am. J. Hum. Genet. 81, 559–575 (2007).

44. H. Cheng, G. T. Concepcion, X. Feng, H. Zhang, H. Li, Haplotype-resolved de novo assembly using phased assembly graphs with hifiasm. Nat. Methods 18, 170–175 (2021).

45. M. J. Roach, S. A. Schmidt, A. R. Borneman, Purge Haplotigs: allelic contig reassignment for third-gen diploid genome assemblies. BMC Bioinformatics 19, 460 (2018).

46. D. R. Laetsch, M. L. Blaxter, BlobTools: Interrogation of genome assemblies. F1000Res. 6, 1287 (2017).

47. C. Camacho, G. Coulouris, V. Avagyan, N. Ma, J. Papadopoulos, K. Bealer, T. L. Madden, BLAST+: architecture and applications. BMC Bioinformatics 10, 421 (2009).

48. E. W. Sayers, E. E. Bolton, J. R. Brister, K. Canese, J. Chan, D. C. Comeau, R. Connor, K. Funk, C. Kelly, S. Kim, T. Madej, A. Marchler-Bauer, C. Lanczycki, S. Lathrop, Z. Lu, F. Thibaud-Nissen, T. Murphy, L. Phan, Y. Skripchenko, T. Tse, J. Wang, R. Williams, B. W. Trawick, K. D. Pruitt, S. T. Sherry, Database resources of the national center for biotechnology information. Nucleic Acids Res. 50, D20–D26 (2022).

49. A. Brandt, P. Tran Van, C. Bluhm, Y. Anselmetti, Z. Dumas, E. Figuet, C. M. François, N. Galtier, B. Heimburger, K. S. Jaron, M. Labédan, M. Maraun, D. J. Parker, M. Robinson-Rechavi, I. Schaefer, P. Simion, S. Scheu, T. Schwander, J. Bast, Haplotype divergence supports long-term asexuality in the oribatid mite Oppiella nova. Proc. Natl. Acad. Sci. U. S. A. 118, e2101485118 (2021).

50. H. Li, R. Durbin, Fast and accurate short read alignment with Burrows-Wheeler transform. Bioinformatics 25, 1754–1760 (2009).

51. C. Zhou, S. A. McCarthy, R. Durbin, YaHS: yet another Hi-C scaffolding tool. Bioinformatics 39 (2023).

52. N. C. Durand, M. S. Shamim, I. Machol, S. S. P. Rao, M. H. Huntley, E. S. Lander, E. L. Aiden, Juicer provides a one-click system for analyzing loop-resolution Hi-C experiments. Cell Syst. 3, 95–98 (2016).

53. J. Wolff, L. Rabbani, R. Gilsbach, G. Richard, T. Manke, R. Backofen, B. A. Grüning, Galaxy HiCExplorer 3: a web server for reproducible Hi-C, capture Hi-C and single-cell Hi-C data analysis, quality control and visualization. Nucleic Acids Res. 48, W177–W184 (2020).

54. M. Manni, M. R. Berkeley, M. Seppey, F. A. Simão, E. M. Zdobnov, BUSCO update: Novel and streamlined workflows along with broader and deeper phylogenetic coverage for scoring of eukaryotic, prokaryotic, and viral genomes. Mol. Biol. Evol. 38, 4647–4654 (2021).

55. D. E. Wood, J. Lu, B. Langmead, Improved metagenomic analysis with Kraken 2. Genome Biol. 20, 257 (2019).

56. S. Capella-Gutiérrez, J. M. Silla-Martínez, T. Gabaldón, trimAl: a tool for automated alignment trimming in large-scale phylogenetic analyses. Bioinformatics 25, 1972–1973 (2009).

57. L.-T. Nguyen, H. A. Schmidt, A. von Haeseler, B. Q. Minh, IQ-TREE: a fast and effective stochastic algorithm for estimating maximum-likelihood phylogenies. Mol. Biol. Evol. 32, 268–274 (2015).

58. H. Shimodaira, M. Hasegawa, Multiple comparisons of log-likelihoods with applications to phylogenetic inference. Mol. Biol. Evol. 16, 1114–1116 (1999).

59. Y. Wang, D. J. Obbard, Experimental estimates of germline mutation rate in eukaryotes: a phylogenetic meta-analysis. Evol. Lett. 7, 216–226 (2023).

60. F. Plazzi, A. Ricci, M. Passamonti, The mitochondrial genome of Bacillus stick insects (Phasmatodea) and the phylogeny of orthopteroid insects. Mol. Phylogenet. Evol. 58, 304–316 (2011).

